# Preserved self-other integration during social decision making among individuals with elevated autistic traits

**DOI:** 10.64898/2026.06.05.728761

**Authors:** Yongling Lin, Elizabeth Pellicano, Cormac Dickson, Nadescha Trudel, MaryAnn Noonan, Patricia Lockwood, Yue-jia Luo, Stephen M Fleming, Marco K Wittmann

## Abstract

Autistic people can find social interactions difficult to navigate, traditionally attributed to difficulties in taking others’ perspectives. However, we have a limited understanding of how autistic people integrate self and other information efficiently during social decision-making. We conducted four highly powered experiments (total N = 1,621) to determine whether autistic traits affect two aspects of self-other integration during social decision making: self-bias and social basis function use. Using Bayesian analyses, we found strong support for the absence of a relationship between autistic traits and either aspect of social decision making, even after controlling for potential confounds (BF_01_ = 32.13 for self-bias, BF_01_ = 7.04 for social basis function use). Our results indicate that variations along autistic traits do not impact how people prioritise self-relevant information (self-bias) or utilize compressed social patterns of interaction (social basis function use) to guide their decisions about oneself and other people. These findings nuance the conceptualisation of social-cognitive processes across autistic traits while highlighting the need for large samples to validate null effects.

## Introduction

Autism is a neurodevelopmental condition that is formally characterised by difficulties in social interaction and communication, along with restricted interests and repetitive behaviours [1]. While there is debate about whether autism is a categorical construct, evidence indicates that autistic traits are expressed in a continuous fashion in the general population [2–5]. Classical theories propose that people with more prominent autistic traits struggle to take other people’s perspectives and see the world from their viewpoint [6–8]. Autistic traits may also affect how people perceive and understand themselves [9–12]. However, social decision making involves more than just understanding oneself or others in isolation; a central challenge is integrating information from both self and others to make good decisions. This problem of integration becomes particularly pressing when information about many individuals needs to be tracked, and when our relationships with them – such as through cooperation and competition – dynamically change [13–15]. To date, we have a limited understanding of how autistic traits impact the integration of self and other information to guide social decision-making in complex environments.

In a social decision-making scenario, we need to track information about others’ identities, such as their attitudes and abilities. However, these pieces of decision information are not integrated linearly - people tend to overweight self-related information relative to information about other people [16–19], known as a “self-bias”. This bias in decision-making may be evolutionarily adaptive, as self-relevant information directly affects survival and may therefore prioritized as more informative [18–21]. However, it is not clear if such self-bias also characterises decision-making in autism. Previous research has come to very different conclusions. Some researchers have argued that autistic people exhibit a lack of self [22–24], while others suggest that the self remains unaltered in autistic people [25–27].

Tracking decision information about every individual becomes computationally demanding as the social group grows. Our recent work shows that the brain encodes combinatorial patterns of how people interact with each other, using these patterns – known as basis functions – as building blocks to compress complex social information and make efficient decisions [28, 29]. Social basis functions encode combinatorial patterns of group relationships, which are particularly useful information in complex social environments [15, 30–32]. Social basis function use is behaviourally characterised by an apparent overreliance on social group structure during decision making: when making decisions about ourselves, our decisions are partly impacted by irrelevant others in line with the social relationships we hold with them [31, 33, 34]. For example, in social interactions with another high-performing individual, people tend to overestimate their own performance when cooperating with them but underestimate their own performance when competing with them [15, 32]. While autism is generally recognized to involve difficulties in understanding others, little research has examined whether autistic traits affect such fundamental representations of social structure, such as social basis functions.

In this work, we investigated whether people varying in autistic traits differ in the integration of self and other information during social decision making. We focused on the above-mentioned two key aspects of self-other integration: self-bias and social basis function use. We used a novel four-player decision-making paradigm to measure self-bias and the use of social basis function [28]. A social structure was established by grouping players into two teams, one comprising the participant (Self, S) and a partner (Partner, P), and the other comprising two other players (Opponents 1 and 2, O1 and O2). On each trial, a sequence of performance cues linked to each player was shown one after another (Fig.1A,B). Participants tracked the cues of each of the four players to make decisions about them afterwards. They were asked to either make a dyadic comparison by recalling the observed performance cues of two players from each team (self and partner decisions) or to compare the two teams overall (group decisions). A crucial design feature of our task was that, from a normative, economic perspective [35], people should attend to all players equally to maximise the points they gain. They were also explicitly instructed that making good decisions on behalf of their partner would increase their points just as much as making good decisions for themselves. Therefore, by comparing decision-making in self and partner decisions, we could distil a precisely controlled measure of self-bias [36–38]. In addition, by statistically assessing the influence of irrelevant players on decisions in line with the social structure, we could measure social basis function use in a precise and quantitative manner [28, 29].

**Fig. 1:**
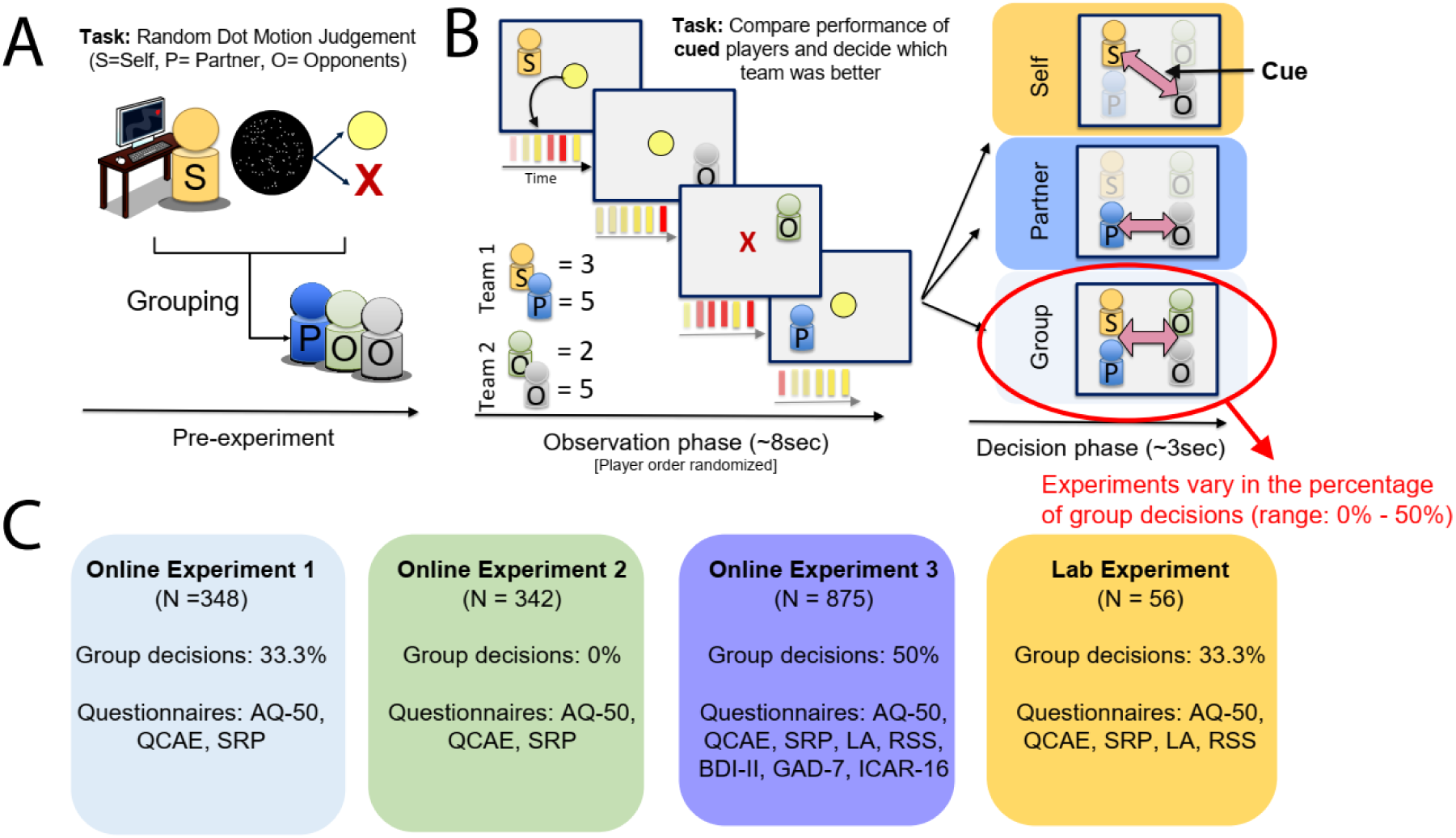
Group decision-making task and study workflow. (**A**). As a pre-experimental task, participants performed a left/right motion judgment task with random dot stimuli, receiving feedback via performance cues: yellow coins (correct) or red crosses (incorrect). Participants were paired with a partner (‘P’) against two opponents (‘O’) before starting the following main task. **(B).** The main task consisted of two phases in each trial: an observation phase and a decision phase. During the observation phase, participants observed the same performance cues as in the pre-experiment for each player unfold over time: Self (S), Partner (P), Opponent 1 (O1), and Opponent 2 (O2) in a random order. Successful performances were shown as yellow coins and erroneous performances were shown as red crosses, mirroring the correctness cues in the pre-experimental task. The participant was asked to aggregate successful performance cues for each player in preparation for the following decision phase. The example trial shows the following scores (= sum of yellow coins per player): an S-score of 3, a P-score of 5, an O1-score of 2, and an O2-score of 5. In the subsequent decision phase, the participant made one of three types of decisions indicated by an arrow cue: Self, partner, and group decisions. The arrow cue indicated the relevant to-be-compared players. Importantly, participants could not predict which players were relevant on a given trial. In self decisions, participants compared their own score with the one from the cued opponent. In partner decisions, they compared their partner’s score with the score of the cued opponent. Finally, in group decisions, they compared the sum of the scores between both groups. **(C).** We performed four experiments: three online experiments (online experiment 1, 2, and 3) and one lab experiment. All experiments followed the same structure, comprising an observation phase and a decision phase. The key differences were the percentage of group decisions, which varied between 0% and 50% (remaining decisions were always in equal part self and partner decisions), and the set of questionnaires used for each experiment (AQ-50: Adult Autism Spectrum Quotient - 50 items; QCAE: Questionnaire of Cognitive and Affective Empathy; SRP: Self-Report Psychopathy Scale; LA: UCLA Loneliness Scale; RSS: Rosenberg Self-Esteem Scale; BDI-II: Beck Depression Inventory-II; GAD-7: Generalized Anxiety Disorder-7-item scale; ICAR-16: International Cognitive Ability Resource).

A key obstacle that limits our understanding is the lack of large-scale data sets to determine the relationship between autistic traits and social decision making [39, 40]. Large sample sizes are particularly crucial for providing strong evidence for null findings [41]. Additionally, we used the Autism Spectrum Quotient questionnaire with 50 items (AQ-50) [42, 43], one of the most frequently used autism assessment tools, to measure autistic trait scores, and to relate them to two characteristic features of social decision making: the self-bias and the use of social basis functions.

To preview our findings, across four experiments totalling 1,621 participants, we show strong evidence for the absence of relationships between autistic trait score and self-bias (BF_01_ = 32.13) or social basis function use (BF_01_ = 7.04) in a general population. The experiments differed in the precise set of questionnaires used, but all experiments included the AQ-50. They also differed in the proportion of group decisions (Fig.1C). We show that the absence of these correlations is not due to an insensitivity of our experimental paradigm to detect the integration of self and other in social decision making: we replicated significant effects of self-bias and social basis function use in all behavioural experiments, whether conducted online or within the laboratory. In addition, we show that our measures of autistic trait score showed expected correlational patterns with other measures, such as empathy questionnaires [44], suggesting our measure of autistic traits had high construct validity. The absence of relationships between autistic trait score and our measures of self-other integration in social decision making remained even after controlling for other psychiatric features and demographics, and even when focusing on socially-specific subscales of the AQ-50. Our findings suggest that people varying in autistic trait scores in the general population exhibit unaltered integration of self and other information regarding self-bias and social basis function use.

## Results

### Measurement of self-bias and social basis function use

We first demonstrate how our experimental paradigm enabled us to measure self-bias and social basis function use, based on the results of Experiment 1. We measured self-bias by comparing the percentage of correct decisions between self and partner decisions. These two decision types occurred in equal numbers in the experiment, were equally important for collecting points in the experiment, and were carefully balanced in terms of decision difficulty. Therefore, any increase in decision accuracy for self compared to partner decisions indicates the prioritization of self-relevant information – a self-bias. Indeed, despite the absence of a task-related incentive, participants made more accurate choices in self decisions compared to partner decisions (*t*(347) = 3.339; *p* < 0.001) (Fig. 2A, B). Note that, while we defined self-bias as a function of decision accuracy, we were also able to replicate this effect in decision reaction times: Participants were significantly faster in self decisions compared to partner decisions across all four experiments we conducted (Supplementary Fig. S1A, B).

**Fig. 2:**
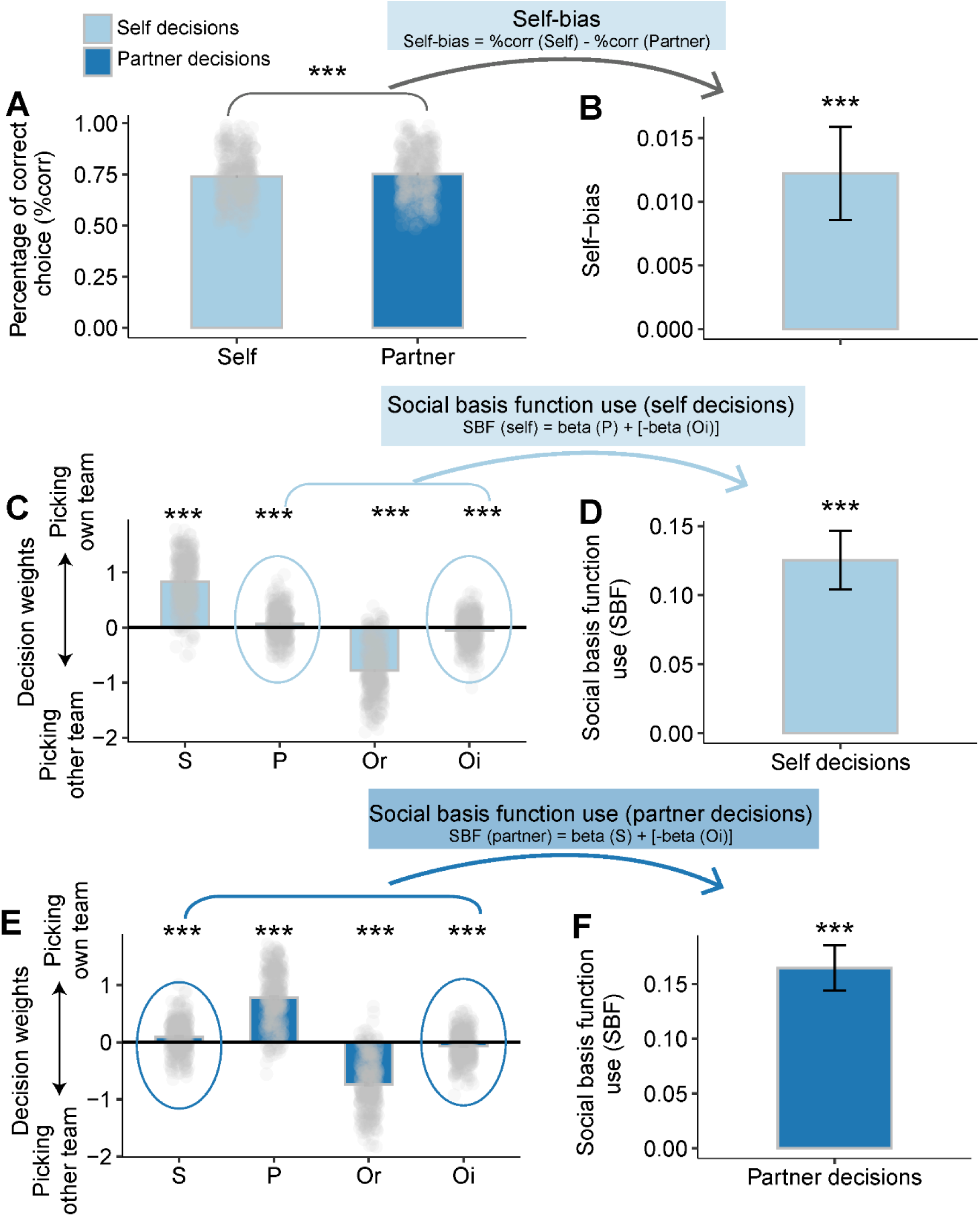
Calculation of self-bias and social basis function use in experiment 1 (n = 348). **(A)** Participants’ decisions were significantly more accurate in self decisions as opposed to partner decisions. **(B)** We calculated self-bias as the difference of the percentage of accurate choices in self decisions and partner decisions (i.e., self-bias = %corr(Self) - %corr(Partner)). **(C)** Effect sizes of logistic GLM predicting self decisions to choose their own team as a function of the performance scores for Self (S), partner (P), relevant opponent (Or) and irrelevant opponent (Oi). We found that the choices were guided by the scores of relevant players (i.e., the self (S) and the relevant opponent (Or)) in self decisions. The positive weight from S-performance indicated that, as they should, participants were increasingly likely to pick their own team if they performed well. At the same time, negative Or-performance effects indicated that as the relevant opponents performed better, participants were less likely to choose their own team, as they should. In addition, we found effects of the irrelevant players (i.e., the partner (P) and the irrelevant opponent (Oi)) consistent with the group membership: participants were more likely to choose their own team when their partner had higher scores, while they were less likely to pick their team if the irrelevant opponent had higher scores. This happened even though the scores of both irrelevant players should be ignored in the decisions. **(D)** We aggregated the effect sizes from irrelevant players (i.e. SBF = beta (P) + [-beta (Oi)] in self decisions) and found a significant social basis function use effect, illustrated in panel D. **(E)** Same regression results in partner decisions. Again, irrelevant players (S and Or in this case) significantly impacted the choice in line with their allegiances with the participant. **(F)** Integrating the effect sizes from irrelevant players (i.e. SBF = beta (S) + [-beta (Oi)] in partner decisions) revealed a significant social basis function use effect. (Error bars are SEM. *** *p* < 0.001.)

To measure the effect of social basis function use, we computed a logistic general linear model (GLM) predicting participants’ trial-by-trial decisions to pick their own team as a function of the performances observed for all four players. We did this separately for self and partner decisions (see Methods). Note that participants were instructed to decide based on the relevant players’ performance alone, for instance, self (S-performance) and relevant opponent (Or-performance) in self decisions. While both effects were strongly present (S-performance: *t*(347) = 34.00; *p* < 0.001, Or-performance: *t*(347) = −31.09; *p* < 0.001), there was clear evidence for additional effects of the partner’s (P) and the irrelevant opponent’s (Oi) performance, even though these players should be ignored in the decision. We observed positive effects of P-performance (*t*(347) = 4.676; *p* < 0.001, and negative effects of Oi-performance (*t*(347) = −3.912; *p* < 0.001) (see Fig. 2C, D). This meant that when the partner scored highly (i.e. many yellow coins were observed for the partner), people were more likely to pick their own team, while a high score of Oi made them more likely to pick the opponent team. We found the same pattern of results in partner decisions. In partner decisions, again, choices were mainly influenced by the relevant players’ scores, which in this case were P-performance (*t*(347) = 30.78; *p* < 0.001) and Or-performance (*t*(347) = −29.60; *p* < 0.001). However, again, irrelevant players’ (i.e., S and Oi) performances significantly impacted decisions in line with their group membership: S-performance exerted a significantly positive effect on choice (*t*(347) = 6.194; *p* < 0.001), and Oi-performance exerted a significantly negative effect (*t*(347) = −5.165; *p* < 0.001) (see Fig. 2E, F).

We quantified the effect of social basis function use (SBF) by integrating the effect sizes of the two irrelevant players in line with their group membership, as in our previous work [15, 30]. As expected, this calculation resulted in significant effects of social basis function use in both self decisions (*t*(347) = 5.904; *p* < 0.001) (see Fig. 2D) and partner decisions (*t*(347) = 8.057; *p* < 0.001) (see Fig. 2F). This demonstrates that our paradigm allowed us to assess self-bias and social basis function use independently.

### Self-bias and effects of social basis function use replicate across experiments and reflect social-cognitive computations

Next, we calculated self-bias and social basis function use as explained in the last section for the remaining experiments. All experiments used the same paradigm, and decision accuracy was comparable across all four experiments (absent main effect: *F*(3,1617) = 0.633, *p* = 0.593, η2 = 0.001, Fig. 3A). Both self-bias and SBF effects replicated across all experiments, suggesting that our paradigm reliably measured both aspects of self-other integration during social decision making. Self-bias was replicated across two other online experiments and one lab-based experiment (*ts* ≥ 2.094; *ps* ≤ 0.041; Fig.3A,B,C). In addition, we replicated these findings using an independent measure of self-bias by quantifying reaction time differences between conditions (Supplementary Fig. S1A,B). Social basis function use also replicated in all studies in both self decisions (*ts* ≥ 3.020; *ps* ≤ 0.003; Fig.3D,E,F) and partner decisions (*ts* ≥ 5.094, *ps* ≤ 0.001; Fig.3G,H,J; see Supplementary Fig. S2 for full GLMs).

**Fig. 3:**
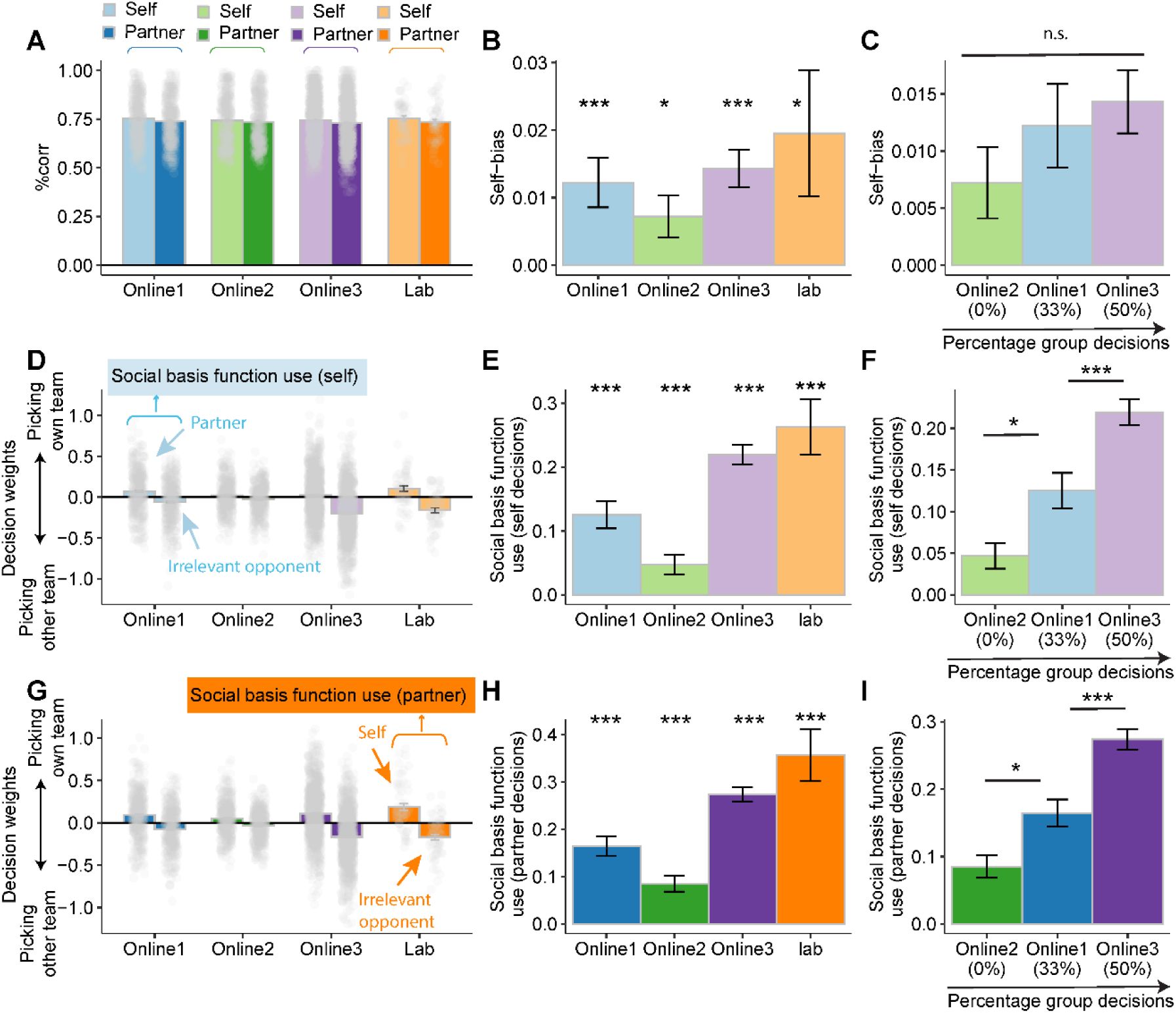
Self-bias and effects of social basis function use replicate across four experiments. We use light colors to indicate self decisions and dark colors for partner decisions. Note that online experiment 1 results were reported in Fig.2 **(A)** Participants showed significantly higher accuracy in self decisions as opposed to partner decisions, i.e. they showed a self-bias. The absence of significant differences in the percentage of correct choices across these four experiments suggests a similar overall level of task difficulty. **(B, C)** The Self-bias was found across all experiments (online experiment 2: *t*(341) = 2.319; *p* = 0.021; online experiment 3: *t*(874) = 5.183; *p* < 0.001), as well as a lab-based experiment (*t*(55) = 2.094; *p* = 0.041). **(D)** Decision weights from the irrelevant players (i.e., partner and uncued opponent) in self decisions across all experiments. **(E)** Social basis function use effects in self decisions were replicated across experiments (online experiment 2: *t*(341) = 3.020; *p* = 0.003; online experiment 3: *t*(874) = 14.183; *p* < 0.001; lab experiment: *t*(55) = 6.047; *p* <0.001). **(F)** Social basis function use effect in self decisions significantly scaled in proportion to the percentage of group decisions. **(G)** Decision weights from the irrelevant players in partner decisions, encompassing self and the uncued opponent. **(H)** Social basis function effects in partner decisions were replicated across three other experiments (online experiment 2: *t*(341) = 5.094; *p* < 0.001; online experiment 3: *t*(874) = 17.897; *p* < 0.001; lab experiment: *t*(55) = 6.499; *p* < 0.001). **(I)** Similar SBF results were found in partner decisions: the effect of social basis function use increased significantly with the percentage of group decisions. (Error bars depict SEM. *** *p* < 0.001, ** *p* < 0.01, * *p* < 0.05.)

We have previously suggested that making decisions on behalf of one’s group might increase social basis function use [45]. As an additional test of whether our measures of self-other integration tracked meaningful social-cognitive computations, we therefore examined the effect of the percentage of group decisions (ranging from 0 to 50%) on social basis function use. To conduct this comparison, we focused on the three online experiments, as they were comparable in all other respects.

As predicted, the effect of social basis function use differed related to the percentage of group decisions (self decisions: *F*(2,1562) = 23.241, *p* < 0.001, η2 = 0.029, Fig. 3F; partner decisions: *F*(2,1562) = 28.745, *p* < 0.001, η2 = 0.035), Fig. 3I). Follow-up post-hoc t-tests revealed that online experiment 2, with 0% of group decisions, showed a lower social basis function use when compared to online experiment 1, with 33% of group decisions (self decisions: *t*(1562) = −2.504, *p* = 0.033; partner decisions: *t*(1562) = −2.548, *p* = 0.029) and which in turn exhibited a lower social basis function effect than online experiment 3, with 50% of group decisions (self decisions: *t*(1562) = - 3.603, *p* < 0.001; partner decisions: *t*(1562) = −4.219, *p* < 0.001). These results suggested that despite receiving similar instructions regarding group structure, group decisions may enhance shared representations in a team setting. These results indicated that simply adopting a group framing may not suffice to foster a sense of group, instead emphasizing the importance of engaging in joint actions to develop overlap between self and others [46–48]. By contrast, as expected, the percentage of group decisions did not significantly modulate self-bias (*F*(2,1562) = 1.123, *p* = 0.326, η2 = 0.001, Fig. 3C).

In addition, to solidify our results even more, we used a second, alternative parameterization of social basis function use. According to our behavioural model, people first identify the group structure and then specify the relevant players within each team [28]. An alternative index of social basis function use can therefore be calculated by considering the time course of how decision variables influence choice on a single trial. This index is characterized by early reliance on the social group structure when people responded quickly. These results were replicated across all four studies, indicating that the task yields well-constructed measures of social basis function use (Supplementary Fig. S3).

### Strong evidence for absent relationships between autistic trait scores and both self-bias and social basis function use

After establishing our behavioural measures of self-other integration, we tested whether autistic trait scores correlated with these measures using Pearson correlations. We used Bayesian analyses to establish evidence in favour of the null hypothesis. BF_01_ indicates Bayes factors in favour of the null, with values larger than 1 indicating increasing evidence for the null compared to the alternative hypothesis. Values larger than 3 indicate moderate evidence, and values larger than 10 indicate strong evidence in favour of the null hypothesis [49, 50].

First, bayesian analyses strongly supported the absence of correlations between self-bias and autistic trait scores across all four experiments (online experiment 1: r = 0.01, *p* = 0.83, BF_01_ = 14.57, Fig. 4A; online experiment 2: r = - 0.04, *p* = 0.42, BF_01_ = 10.65, Fig. 4B; online experiment 3: r = 0.008, *p* = 0.81, BF_01_ = 22.95, Fig. 4C; lab experiment: r = 0.012, *p* = 0.93, BF_01_ = 5.98, Fig. 4D). The absence of a correlation was consistently replicated when examing the relationship between autistic trait scores and self-bias in reaction time – an alternative, independent measure of self-bias (BF_01_ = 27.27 in the combined dataset, Supplementary Fig. S1C).

**Fig. 4:**
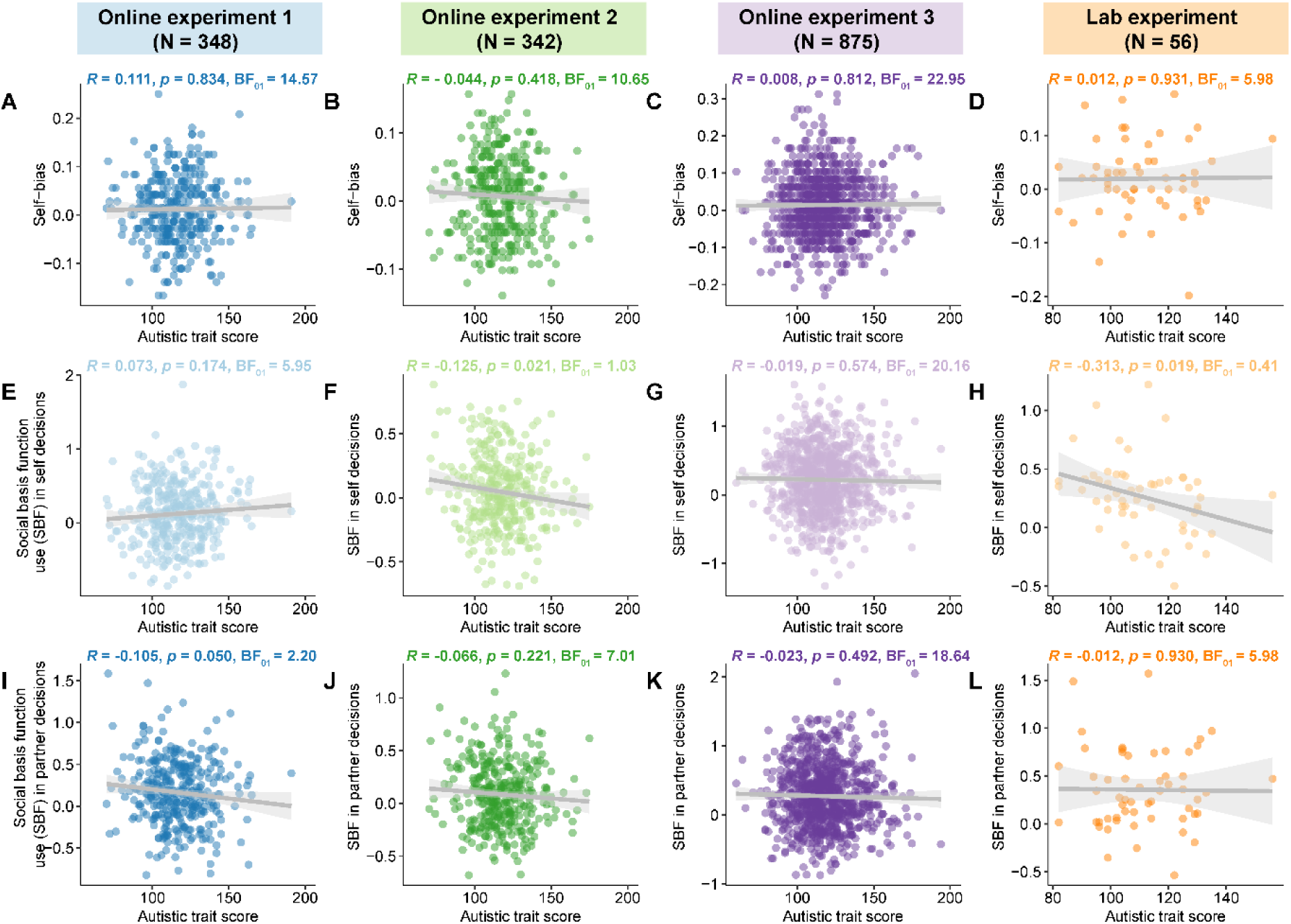
Evidence for an absent relationship between autistic trait scores and the integration of self and other. **(A, B, C, D)** Bayesian analyses showed evidence for an absent relationship between self-bias and autistic trait scores across four experiments. **(E, F, G, H)** An absent correlation was observed between autistic trait score and social basis function use in self decisions in online experiments 1 (n = 348) and 3 (n = 875), with Bayesian analysis strongly supporting the null. In both online experiment 2 (n = 342) and lab experiment (n=56), autistic trait scores showed a slight negative correlation with social basis function use effect in self decisions. However, Bayesian analysis provided weak evidence for the alternative hypothesis in the lab experiment and no supporting evidence in online experiment 2. **(I, J, K, L)** We found an absent correlation between autistic trait score and social basis function usein partner decisions across online experiments 2, 3, and the lab experiment. While online experiment 1 showed a marginally significant correlation, Bayesian analysis still supported the null hypothesis. Overall, there was no consistent and conclusive correlation between autistic trait score and social basis function use effect in either self or partner decisions.

Second, we examined whether autistic trait scores correlated with the effects of social basis function use in self or partner decisions. We found evidence for absent correlations between autistic trait scores and the effect of social basis function use in both self decisions (online experiment 1: r = 0.073, *p* = 0.174, BF_01_ = 5.95, Fig. 4E; online experiment 2: r = - 0.125, *p* = 0.021, BF_01_ = 1.03, Fig. 4F; online experiment 3: r = - 0.019, *p* = 0.574, BF_01_ = 20.16, Fig. 4G; lab experiment: r = - 0.313, *p* = 0.019, BF_01_ = 0.41, Fig. 4H) and partner decisions (online experiment 1: r = - 0.105, *p* = 0.050, BF_01_ = 2.20, Fig. 4I; online experiment 2: r = - 0.066, *p* = 0.221, BF_01_ = 7.01, Fig. 4J; online experiment 3: r = - 0.023, *p* = 0.492, BF_01_ = 18.64, Fig. 4K, lab experiment: r = - 0.012, *p* = 0.930, BF_01_ = 5.98, Fig. 4L). Note that some correlations do reach individual significance using classical statistical methods. However, even in cases of significant correlations between autistic trait scores and social basis function use, these results are still characterised by greater evidence for the null than the alternative hypothesis (online experiment 1: partner decisions, online experiment 2: self decisions). The only exception was the lab experiment, where the correlation between autistic trait scores and social basis function use for self decisions weakly supported the alternative hypothesis (BF_01_ = 0.41; Fig. 4H). But even then, this result would not survive correction for multiple comparisons, is only observed in the study with the smallest sample size, and needs to be interpreted against a larger pattern of null correlations. Note that in the online experiment 3 with the largest sample size (N = 875), no significant correlation was found, with overwhelming evidence for the null (BF_01_ > 18 in both cases, reflecting “strong” evidence for the null) [49, 51]. These results remained robust, with marginal significant correlations from online experiments 1 and 2 becoming non-significant when analysed using non-parametric Kendall’s tau correlation (see Supplementary Fig. S4). We then collapsed the data across all experiments and examined the composite correlations between autistic trait scores and our two markers of self-other integration. Overall, there was strong evidence for the absence of a relationship between autistic trait scores and self-bias (r < - 0.001, BF_01_ = 32.13), and, to a lesser degree, also for social basis function use (self decisions: r = - 0.021; BF_01_ = 22.55; partner decisions: r = - 0.046; BF_01_ = 5.88; overall social basis function use, calculated based on the mean values over self and partner decisions: r = - 0.043; BF_01_ = 7.04).

Lastly, we examined whether autistic trait scores correlated with an alternative behavioural signature of social basis function use, quantified by the temporal priorisation of social group structure during decision making (Supplementary Fig. S3). Again, Bayesian analyses strongly supported the absence of a correlation between this index and autistic trait scores (In combined dataset, self decisions: BF_01_ = 27.28; partner decisions: BF_01_ = 31.04, Supplementary Fig 3E, J).

### The integration of self and other remains unaltered across AQ scoring variations, subscales, and after controlling for confounds

An absence of a relationship between self-other integration and autistic trait scores could be caused by not measuring one or the other properly. We have shown that our task reliably measured self-bias and social basis function use, and that the strengths of social basis function use covaried with subtle changes in our experimental paradigm (Fig.3). In addition, we now show that our measurements of autistic trait score captured component aspects of the relevant construct: autistic trait scores repeatedly and consistently showed the expected patterns of correlation with empathy and other mental health measures (Supplementary Fig. S5).

Next, we focused specifically on socially relevant subscales of the AQ questionnaire and also considered alternative scoring methods of the questionnaire. We first investigated whether self and other integration correlates with any of the five original AQ subscales, including social skill, attention switching, attention to detail, communication, and imagination [42]. Moreover, we also examined its relationship with two- and three-factor structure models of AQ subscales, which have been well-established in previous research and shown to improve internal consistency [52–54].

The resulting 27 Pearson correlations were statistically non-significant (rs ≤ |0.05|, *ps* ≥ 0.115), except that the correlation between AQ-Communication subscale and social basis function use in partner decisions narrowly reached significance (r = - 0.053, *p* = 0.034, Fig. 5A). However, this did not survive correction for multiple comparisons.

**Fig. 5:**
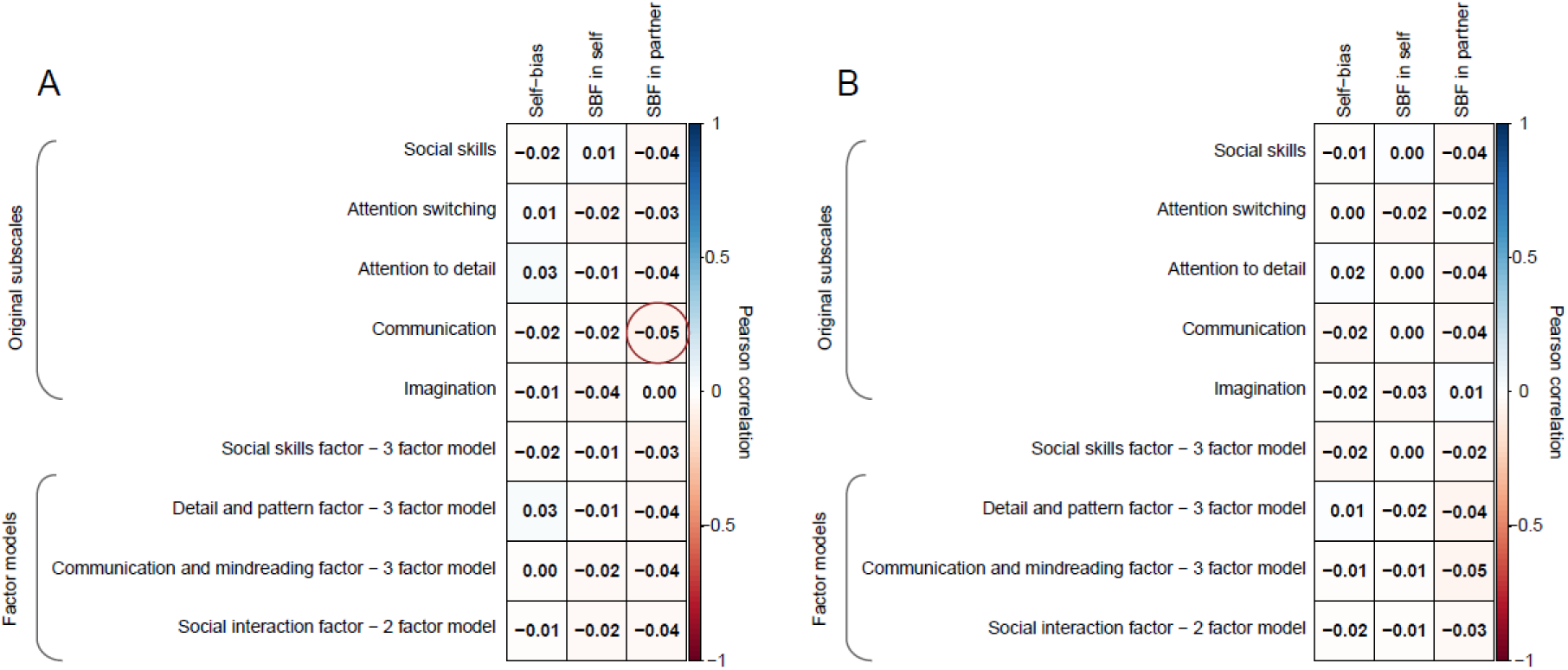
Effects of interest do not change with scoring variations of the AQ. **(A)** We examined Pearson correlations between three markers of self-other integration (self-bias, social basis function use in self decisions, social basis function use in partner decisions) and subscales of autistic trait scores. We collapsed our behavioural data across all four experiments. The AQ subscales comprised not only the five original subscales (social skills, attention switching, attention to detail, communication, and imagination), but also the further established 2- or 3-factor models of AQ-50 autistic trait scores. In the 3-factor model, the three factors are the social skills factor, the detail and patterns factor, and the communication and mindreading factor. In the 2-factor model, the factors are the original subscale of attention to detail as one factor, and a social interaction factor comprising the remaining four original subscales. The absolute values of correlation coefficients were all smaller than 0.05. The only significant correlation (*p* = 0.034) between social basis function use in partner decisions and AQ-communication scores did not survive correction for multiple comparisons. **(B)** We replicated the correlations using the binary AQ scoring system, categorizing items as autistic (1) and non-autistic (0). All non-significant findings remained consistent. Note that the one significant relationship identified between the AQ-communication subscale and social basis function use in partner decisions in panel **A** became non-significant in the corresponding binary-scored AQ subscale analysis in panel **B**.

We also considered the impact of our four-point Likert scoring method used to calculate autistic scores, which has previously been found to improve item discriminability [52, 55]. However, for completeness, we also reanalysed our data using the original binary scoring method, categorizing items as autistic (1) and non-autistic (0), resulting in an overall score range of 0-50 [42]. We repeated our above approach of calculating subscales within the AQ. None of the 27 calculated correlations reached significance (rs ≤ |0.05|, *p* ≥ 0.063, Fig. 5B), and even the correlation observed in Fig.5A was now insignificant.

We further divided participants into two groups based on the established clinical threshold (AQ score ≥ 32, using the binary scoring system) from the previous studies [42, 43, 56]. This grouping allowed us to examine if participants above the threshold showed altered self-other integration compared to those below it. Our results showed that participants scoring above the threshold showed a comparable pattern across these integration measures to those below the threshold (independent samples t-tests: self-bias: BF_01_ = 8.83; social basis function use in self decisions: BF_01_ = 7.19; social basis function use in partner decisions: BF_01_ = 11.84) (see Supplementary Fig. S6).

Additionally, we explored whether controlling for demographic and mental health questionnaire variables would reveal the potential relationships between autistic trait scores and the behavioural markers of self-other integration. General linear models showed no evidence that autistic trait scores predicted self-bias or social basis function use after accounting for potential confounds, even when testing for ‘U-shaped’ relationships using squared scores (Supplementary Fig. S7). These additional analyses further support that our results reflect a genuine absence of a relationship.

## Discussion

Autistic people can find it difficult to understand others in social scenarios. However, while much research has focused on social perspective taking in autism [6–8], less attention has been paid to how people with autistic traits integrate information about themselves and others during social decision making – a process that is crucial for navigating social situations. Our study aimed at investigating whether and how individual differences in autistic traits impact self-other integration when making social decisions. Across four highly powered experiments with 1,621 participants, we examined whether autistic trait scores relate to two aspects of self-other integration: self-bias [17–19] and social basis function use [15, 28]. We found evidence for the absence of a relationship between autistic trait scores and these markers of integration during social decision making, even after controlling for confounds or analyzing socially-specific subscales of the AQ-50 [42]. On aggregate, Bayasian analyses provided strong evidence for null results [57].

A self-bias is characterised by prioritizing self-relevant information over other information when making social decisions [18, 21, 28]. Self-biases are also commonly observed in perception and memory domains, such as faster recognition of one’s own face [17, 58] or quicker judgments when rating trait terms in relation to oneself [59]. Across our four studies, participants were consistently more accurate when deciding on behalf of themselves compared to deciding on behalf of a social partner – even though the financial incentive was the same in both cases. A self-bias in decision making may be evolutionarily adaptive because self-relevant information is more closely related to one’s own survival [17, 18, 60]. Our results indicate that such priorization of self-related information during social decision-making is equally common among people with autistic traits: they similarly made more accurate and quicker decisions when deciding about themselves. The large sample size of the present study strengthens this conclusion and helps disambiguate previously conflicting findings in the literature on self-bias in autistic people [22–27].

When integrating different pieces of social information during social decision making, we not only process information differently based on social identity, but instead we also rely on our prior knowledge of social relationships [61–64]. We have suggested that basis functions are used to compress information along the principal social relationships that exist in a group [28, 29]. This abstract coding scheme is particularly useful as computational demands increase because it allows efficient compression of social information. As previously, in this study, we measured social basis function use as a behavioural index of the over-reliance on inherent group structure during social decision-making [15, 28, 30]. We consistently replicated such over-reliance over our four studies and found that this effect is modulated by the fraction of decisions that are made on behalf of the whole group together. Using the same index, we have in the past shown social basis function use increases during adolescence [29]. By contrast, here, we find that social basis function use remains preserved among individuals with autistic traits, suggesting that people with autistic traits integrate information about this aspect of social structure similarly to other people during social decision making. While specific social-cognitive processes such as perspective-taking may be changed in autism [6, 8, 12], our findings suggest that the ability to use compressed representations for navigating complex social decisions remains unaffected among individuals with autistic traits.

Together, our results suggest that social decision making – in terms of self-bias and social basis function use - is preserved among individuals with varying autistic traits. This raises the question: if not self–other integration, how else might autistic traits impact social decision making? One plausible target is *social inference*. Inference relates to the prediction of latent states based on observable data and an internal model [65]. Our task explicitly probes how people integrate identities and relationship structure during social decision making when those quantities are observed, but it does not require them to infer hidden variables or to update beliefs about changing social dynamics. This suggestion is in line with the evidence on social perspective-taking cited above [6, 8, 12], and is also consistent with recent results suggesting that people with autistic traits may struggle with inferring goals of others during a gambling task [66] and that they show a reduced ability to predict other people’s choices in a mentalising task [67]. Other recent work found that autistic people showed a larger social distance during a social navigation task, while, consistent with our finding, the directional relationship mappings (such as power and affiliation dimensions) in geometric space did not show significant differences from comparison groups [68]. This suggests that at least certain aspects of social decision-making may be preserved. Our results open the door to future work examining higher-order aspects of social decision-making with the goal of narrowing down how autistic traits affect the full breadth of social-cognitive processes.

Since we are arguing for the absence of an effect of autistic traits, it is important to consider whether our design is sufficiently protected against false negatives. We think these are very unlikely for the following reasons. First, our sample size is substantially larger than the samples used in previous work to examine this question [25–27, 36, 69, 70], increasing the likelihood of identifying an effect if present. Second, we consistently replicated self-bias and social basis function use in four experiments, demonstrating that our task captures social-cognitive processes well, including the subtle scaling of social basis function use with the percentage of group decisions (Fig.3). Our task is comparable to other tasks routinely used in social psychology research [15, 71, 72] including those which have identified relationships between autistic traits and individual difference markers [67]. Therefore, our task appears to be well suited to identify relationships between self-other integration and autistic traits, should they exist. Third, we conducted sensitivity analyses using different measurements of autistic traits (AQ scoring variations and social-specific subscales) and controlled for potential confounds to rule out effects which may have been masked by measurement artifacts. No significant effects emerged even after accounting for all these variations. Finally, one might argue that online experiments may invoke weaker social engagement than lab studies [73–75], potentially limiting our ability to detect true effects. However, both questionnaire and decision-making measures from online participants aligned with those obtained in an in-person, laboratory sample. These findings indicate that online studies effectively captured the social task dynamics. Together, these considerations emphasize the importance of large samples to establish meaningful null effects in computational psychiatry.

Our study examined autistic trait scores in the general population rather than people with a diagnosis of autism. Although the AQ-50 shows good sensitivity and specificity in clinical samples [52, 56], diagnosed autistic people typically show higher autistic trait scores. Our study found no differences in the integration of self and other information across the general population, even when we looked specifically at people who scored above the established questionnaire cutoff point associated with an autism diagnosis. However, scoring above this cutoff is not equivalent to receiving a clinical diagnosis, and there might still be differences between clinically-diagnosed and trait-defined autism [76]. Therefore, we should be cautious when generalizing the current findings to diagnosed autistic people [77]. This highlights the need for further studies examining self-other integration across the entire autism spectrum, particularly including autistic people with clinical diagnoses.

In summary, across four experiments totalling 1,621 participants, we found strong evidence for the absence of a relationship between autistic trait scores and markers of self-other integration: self-bias and social basis function use. Our findings suggest that people with varying autistic trait scores exhibit a comparable prioritisation of self-relevant information and share similar tendencies to compress social information using combinatorial coding of group relationships within a social group. These findings help further refine our understanding of the social capabilities of autistic people, which include unaltered computational processes supporting self-other integration. They also highlight the importance of conducting replication studies and employing large sample sizes in the field of computational psychiatry.

## Methods

### Participants

We conducted four experiments (three online experiments and one laboratory experiment) to examine the relationship between autistic traits and self-other integration. Individuals of all genders, including female, male, and non-binary, in the United States and the United Kingdom were eligible to participate. All participants were required to be within the age range of 18-40 and have fluency in English. Online participants received £9 per hour for participating in the experiment. All online data was collected via Prolific (https://www.prolific.co/) [78], with pre-defined exclusion criteria applied to retain only valid responses (Supplementary Table S1). All experimental methods and procedures were approved by the ethics committee of the University of Oxford, and all participants provided informed consent (online experiments: MSD reference number: R70000/RE001; lab experiment: MSD reference number: R60547/RE001). Different participants took part in each experiment.

- **Online experiment 1.** We collected data from 404 participants. The exclusion criteria resulted in the exclusion of 56 participants (13.86%) from further analysis, leaving a final sample of 348 participants (145 females, 201 males, and 2 non-binary; median age = 31).
- **Online experiment 2.** We collected data from 402 participants; 60 participants (14.93%) were disqualified, leaving a final sample of 342 participants (185 females, 152 males, and 5 non-binary; median age = 31).
- **Online experiment 3.** A total of 1050 participants were collected, and 175 participants (16.67%) were excluded after applying the exclusion criteria, leaving 875 participants for analysis (409 females, 454 males, and 12 non-binary; median age = 30).
- **Lab experiment.** We recruited 59 participants in the lab experiment. Two participants did not complete the whole experiment, and one participant repeatedly fell asleep during the experiment. The final sample included 56 participants (33 females, 23 males, 0 non-binary; median age = 23). Participants received £50 for participating in the experiment, plus extra earnings which were allocated according to their task performance. The high payment was due to their involvement in functional magnetic resonance imaging (fMRI) recording, which is unrelated to the current study and will be reported separately in our other studies [28].

After the task instructions (i.e., after ∼7 minutes of participation) in the three online experiments, participants were asked to respond to a comprehension check comprising multiple-choice questions that required an understanding of the task rules. The study would be aborted if they incorrectly responded to any of the three questions in the comprehension check. Excluded participants would be reimbursed for their time. For those participants who have taken part in our experiment multiple times, we only considered their initial completed dataset and discarded subsequent datasets. Therefore, only datasets from participants who passed the comprehension check and completed the study on their initial attempt were retained. Following that, we excluded inattentive participants based on pre-defined exclusion criteria aligned with standard guidelines for online data collection [79]. The criteria and the number of excluded participants were reported in Supplementary Table S1. Our exclusion rate falls within the range reported in a recent meta-analysis, which found that between 3% and 37% of the sample is excluded in online data collection [80]. All these behavioural data were previously reported in [28], but with a fundamentally different aim. Previously, we focused on the basis functions that structure social interactions in dorsomedial prefrontal cortex. Now, questions about the neural mechanisms that people use to summarise social environments are no longer of primary importance. Instead, the current work focuses on individual differences in questionnaire-assessed measures of autistic traits and mental health, and their relationship to markers of self-other integration. The questionnaire data presented in the current study were not part of the previous publication.

### Experimental procedures

Across all four experiments, participants first completed a pre-experiment phase, followed by a multi-person group decision-making task and self-reported questionnaires. For the three online experiments, the task was developed using jspsych [81], where participants accessed a web link to begin and conclude the experiment. The laboratory experiment was programmed in MATLAB using Psychtoolbox-3 (http://psychtoolbox.org).

The pre-experiment involved a random dot kinematogram (RDK) judgement task, where the participant pressed left/right buttons to indicate leftwards/rightwards motion directions of the RDK stimuli. Importantly, performance cues were used to indicate correct and incorrect RDK performance (a yellow coin and a red ‘X’). These performance cues were the same also used in the subsequent group decision-making task, allowing participants to familiarise themselves with these performance cues before the main task. The pre-experiment took approximately 3 minutes in the online experiments and approximately 25 minutes in the laboratory experiment, where more time could be allocated.

Afterwards, the group decision-making task took place. It involved four players (as in [28]). The players were divided into two teams: the participant and their partner (“P”) formed one team, and the other two players formed the opponent’s team (“O1”, “O2”). Each trial comprised an observation phase and a decision phase. In the observation phase, performance cues relating to all players were shown, and in the decision phase, participants made decisions about the presented performance cues from memory.

During the observation phase, each of the four players’ scores was represented by a sequence of six performance cues, presented centrally on the screen. These performance cues were identical to those used during the pre-experiment, with yellow ‘coins’ indicating successful performances and red ‘X’ cues indicating erroneous performances. While the performance cues were presented rapidly (300 ms per cue, with 100 ms inter-stimulus intervals), a moving RDK was displayed at the location of the relevant player to indicate which player the performances referred to. Note that in the main experiment, no RDK judgements were made. Participants were simply asked to aggregate the performance cues towards an overall score per player. To determine each player’s score, participants needed to keep track of the series of six performance cues, accumulating the yellow ‘success’ cues while disregarding the red ‘error’ cues. The number of successful performances, which range between 0 and 6, reflected the resulting scores of each player. The performance cues for all four players were displayed in a random but counterbalanced sequence.

On each trial, after the observation phase, the decision phase followed. During the decision phase, participants retrospectively compared performance scores between players from their own team and the opposing team, using an arrow cue to indicate the players to compare. The decision was to choose the team that performed better, with each decision phase comprising two decisions based on the same set of performances in the trial. Each decision lasted until a response was given. Afterwards, a box appeared around the team that the participant had picked for 0.5 seconds. Note that temporal jitters were applied to the lab-based experiment, given that fMRI data were collected simultaneously [28].

Three decision types were employed in the decision phase: self decisions, partner decisions, group decisions. Decisions occurred pseudorandomly and could not be predicted in advance. The decision type was indicated by an arrow that appeared and pointed towards decision-relevant players. In self decisions, the participant’s performance was compared with the performance of one of the two opponents. The performance of the other two players had to be ignored. In partner decisions, the partner’s performance was compared with the performance of one of the two opponents, ignoring the performances of self and the other irrelevant opponent. Finally, in group decisions, the sum of the performances of both groups was compared. In addition, participants had to factor in a non-social bonus, indicated by the color of the arrow cue with a value of 0.5, which determined whether to add points to one’s own team’s performance (yellow arrow) or to the opponent team’s performance (red arrow). The bonus was present on every decision, and this meant that the decision always had a correct response, even if the performance scores of the cued players were identical. The laboratory experiment also comprised a bonus on every decision, but with a wider range of bonus values (0.5 or 1.5). The bonus was displayed as yellow and red coins at the top of the decision arrow, indicating whether points would be added to one’s own team’s performance (yellow coin) or to the opponent team’s performance (red coin). Participants’ incentive to do the task was to collect ‘points’. This worked in the following way. If participants chose the player from their own team, then the points gained or lost were equal to the performance difference between the two cued players. If the one from their own team was indeed better, they received points equivalent to the true performance difference. Otherwise, they lost the points equivalent to the true performance difference. The outcome of choosing the player from the opponent team always led to a payoff of zero.

Well-defined input schedules were used for participants. Slightly different schedules were applied across the online experiments, including variations in the percentage of group decisions (Online experiment 1: 33% of group decisions; Online experiment 2: 0% of group decisions; Online experiment 3: 50% of group decisions; Lab experiment: 33% of group decisions) and trial numbers (Online experiment 1: 108 trials; Online experiment 2: 108 trials; Online experiment 3: 96 trials; Lab experiment: 144 trials.). For the lab experiment, participant-specific schedules were created. More information on these schedules can be found in [28].

### Self-report questionnaires

Across all four experiments, all participants completed the 50-item Adult Autism Spectrum Quotient, which is a widely-used self-report measure of autistic traits [82]. They also completed the Questionnaire of Cognitive and Affective Empathy [44] and Self-Report Psychopathy Scale [83]. In online experiment 3 and lab experiment, we expanded our questionnaire set to include additional mental health questionnaires. Both experiments included the Rosenberg Self-Esteem Scale [84] and the UCLA Loneliness Scale [85]. In addition, online experiment 3 also included the Beck Depression Inventory–II [86]; Generalized Anxiety Disorder 7-item scale [87]; International Cognitive Ability Resource [88].

The full set of questionnaires used was the following:

- The 50-item Adult Autism Spectrum Quotient (AQ-50) [82]. The AQ consists of 50 items, each of which is scored on a scale from one to four. The items were reversed where necessary. The total AQ score was calculated using the four-point Likert scoring method and reported in the main text. This Likert scoring retains valuable scale information to discriminate between individuals [52, 70]. For completeness, the autistic trait score was also calculated using a binary scoring system (1 for autistic, 0 for non-autistic), and reported values were applicable.
- Self-Report Psychopathy Scale (SRP) [83]. SRP is a 29-item scale that measures psychopathic traits in noninstitutionalized samples. Each SRP item uses a five-point Likert scale and some item scores were reversed when necessary. All SRP items are summed to produce the total psychopathic scores.
- Questionnaire of Cognitive and Affective Empathy (QCAE) [44]. QCAE is a 31-item scale, and each of the items is rated on the level of agreement using a four-point scale from “strongly disagree” to “strongly agree”. The item scores are reversed where necessary. We calculated the scores on the cognitive empathy subscales and the scores on the affective empathy subscales. The sum of cognitive and affective empathy scores produced the total empathy score.

We included the following additional questionnaires in online experiment 3, with the first two questionnaires also being included in the laboratory experiment.

- Rosenberg Self-Esteem Scale (RSS) [84]. RSS is a 10-item scale. Each item is rated in a four-point Likert scale format and some items were reversed scored if necessary. Summary scores on RSS were calculated.
- UCLA Loneliness Scale (LA) [85]. LA is a 20-item scale. Similar with RSS, each item is rated in the four-point Likert scale format and some items were reversed if necessary. summary scores on LA were calculated.
- Beck Depression Inventory-II (BDI-II) [86] . BDI-II is a 21-items scale. Each item is rated on a four-point Likert scale ranging from zero to three. No reverse scoring was required. Summary scores for BDI-II were calculated.
- Generalized Anxiety Disorder 7-item scale (GAD-7) [87]. GAD-7 is a 7-item scale. Each item is rated on a four-point scale, ranging from ‘Not at all’ (0) to ‘Nearly every day’ (3). No reverse scoring was required. The total score of GAD-7 ranges from 0 to 21.
- International Cognitive Ability Resource (ICAR-16) [88]. ICAR is a 16-items scale. Summary scores were calculated based on the number of items answered correctly, assigning a score of 1 for correct responses and 0 for incorrect responses.

### Statistical Analysis

We analysed and visualised data using MATLAB 2021b, JASP version 0.18, and R version 4.1.3. Two-tailed tests were used for all analyses. We reported *p* values at a 0.05 alpha level (two-sided) of the test statistic. The results are reported as Mean ± SE. The questionnaire responses are aggregated by calculating their summary scores.

#### Self-bias

We first calculated the percentage of correct decisions and median reaction times separately for self and partner decisions. We conducted a paired-wise t-test to compare the percentage of correct decisions in self and partner decisions. We defined self-bias by calculating the difference in the percentage of correct choice between self decisions and partner decisions (see Eq.1).

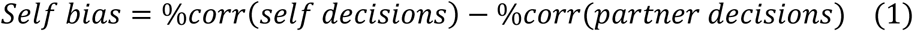

Note that %corr is the percentage of correct decisions in each decision type.

#### Social basis function use in dyadic decisions

We applied a logistic GLM separately for self and partner decisions to predict whether participants picked their own team as the higher performer on the current decision. The dependent variable is the decision that one’s own team member performed better (1 = yes, 0 = no). All regressors were normalised (a mean of zero and a standard deviation of 1). We included the following regressors: the performances from four players (i.e. a number between 0 and 6 for each player on each trial; equivalent to the number of positive performance cues) and the bonus on the current decision (see Eq.2).

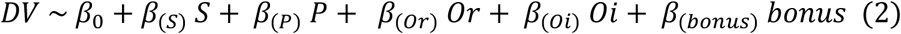

Note that S, P, Or, Oi refers to the performance scores from self, partner, the relevant opponent (i.e., the opponent that self or partner were compared with on the current decision, Or) and the irrelevant opponent (i.e., the opponent were not compared with on the current decision, Oi). The ‘bonus’ refers to the assigned bonus on the current decision (worth either −0.5 or +0.5 in online experiments, or worth 1.5, −0.5, +0.5, or 1.5 in the lab experiment). The dependent variable (DV) is whether participants picked their own team on the current decision (1 = own team, 0 = opponent team).

Ridge regressions were used to estimate the regressors’ beta weights [89–91], which penalizes large beta weights according to a regularization coefficient *λ* and thus prevents overfitting and improves generalization. This is appropriate for cases such as ours when there are many regressors and comparatively few trials. We applied the regression model using MATLAB’s lassoglm (setting Alpha to a very small value) in the following way. To determine an appropriate regularization coefficient *λ*, the GLM was repeatedly fitted to each individual data set with *λ* varied between zero to 10^−3^ to 10^−1^ (log-spaced). At each fit, a three-fold cross-validation approach was used to determine the overall model deviance for each *λ* for all combined data sets. We repeated this procedure twice. Finally, the *λ* that resulted in the lowest overall model deviance was selected. This is the *λ* with the best cross-validated model fit, which was then used to run the ridge GLM of interest. Critically, whenever relevant, the same best-fitting *λ* was used for all regression models of all participants, irrespective of condition, to ensure fair statistical comparisons of beta weights within and across conditions and across four experiments.

The use of social basis function (SBF) was defined in reference to the sum-up effect sizes obtained in the ridge regressions for the irrelevant players in dyadic decisions (i.e., P and Oi in self decisions; S and Oi in partner decisions). Before summing up, we inverted the beta value associated with the irrelevant opponent to account for its negative impact on choice (e.g., SBF(self-decisions) = beta(P) + [-beta(Oi)]). We aggregated the effects from both irrelevant players in the following way for self decisions and for partner decisions separately (See Eq. 3).

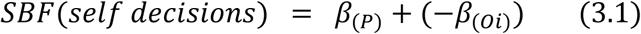

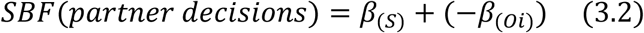

Note the sign of Oi is inverted to account for the fact that in social basis function use, collaborators are weighted positively in accordance with one’s own team, and competitors are weighted negatively in accordance with the opponents’ team [15, 30].

#### Correlating questionnaire scores with self-other integrating effects

We used two correlation measures. Pearson correlations were used and reported in the main text, while the results were reported using Kendall’s rank methods in the supplementary materials. Note that using different correlation methods did not yield any conflicting results. Fisher’s Z-transformation of correlation coefficients was used to examine significant differences between correlation coefficients. To quantify the evidence for the null hypotheses compared to the alternative hypothesis, we implemented a Bayesian inferential approach using JASP and calculated the Bayes factor in favour of the null (BF_01_). In brief, the BF_01_ is the ratio between the marginal likelihoods of the null model (H0) and the alternative model (H1), representing the degree of confidence that both distributions are indeed different [51]. A value of 1 in BF_01_ indicates that H0 and H1 are equally likely given the data, and values above 1 indicate greater support for H0. For instance, ‘BF_01_ = 5’ means that the given data are 5 times more likely under null model (H0) than under the alternative model (H1). By convention, BF_01_ between 1 and 3 is considered weak evidence in favour of the null model, a Bayes factor between 3 and 10 is considered moderate evidence, and a Bayes factor greater than 10 is considered strong evidence. We used a default prior option, a Cauchy distribution with spread r set to 1/√2.

## Acknowledgements

This project was funded by the UCL Institute of Mental Health and the China Scholarship Council (Grant No: 202206040126). We thank Anthony David for his helpful comments on the paper. MKW was funded by an UK Research and Innovation guarantee grant (UKRI; under the UK government’s Horizon Europe funding guarantee UKRI336 [selected as ERC Starting Grant, EC reference 1011159]) and a Medical Research Council grant (MRC; MR/Y010477/1).

## Data availability

All data, analysis scripts, and computational environment are available at https://github.com/YonglingLin/autistic2023.

## Supplemental Materials

### Supplementary tables

**Supplementary Table S1.** Description of online sample in relation to exclusion criteria. We used the following exclusion criteria for the online data. *Exclusion 1*: Participants who took longer than 2 hours to complete the task or the questionnaires were excluded from data analysis; *Exclusion 2*: Participants who showed a choice repetition bias in the study were excluded from data analysis. A choice repetition bias was defined as picking the same choice (left or right button) in more than 85% of trials of either the pre-experiment (see below for more details) or the main task; *Exclusion 3*: Participants who performed at below- or near-chance in the main task (judged as < 55% correct decisions; chance performance was 50%) were excluded from data analysis; *Exclusion 4*: Participants who took longer than 30 seconds to make decisions in more than 10% of trials in the main experiment were excluded from data analysis; *Exclusion 5*: Participants who responded incorrectly to one or more of the three ‘catch’ questions embedded in the questionnaires were excluded from data analysis. These catch questions had the following structure: ‘If you are paying attention to these questions, please select x as your answer’. Excluded participants per criterion are described in absolute numbers and percent of the original sample (indicated in brackets).

|  | Exclusion 1 N(%) | Exclusion 2 N(%) | Exclusion 3 N(%) | Exclusion 4 N(%) | Exclusion 5 N(%) | Remaining participants |
| --- | --- | --- | --- | --- | --- | --- |
| Online Experiment 1 (N = 404) | 0 (0%) | 8 (1.98%) | 32 (7.92%) | 0 (0%) | 19 (4.70%) | 348 (86.14%) |
| Online Experiment 2 (N = 402) | 0 (0%) | 3 (0.75%) | 46 (11.44%) | 0 (0%) | 17 (4.23%) | 342 (85.07%) |
| Online Experiment 3 (N = 1015) | 9 (0.89%) | 14 (1.38%) | 135 (13.30%) | 6 (0.59%) | 41 (4.04%) | 875 (83.33%) |

Note: the percentage of exclusions includes some participants who violated more than one exclusion criterion.

### Supplementary figures

**Supplementary Figure S1.**
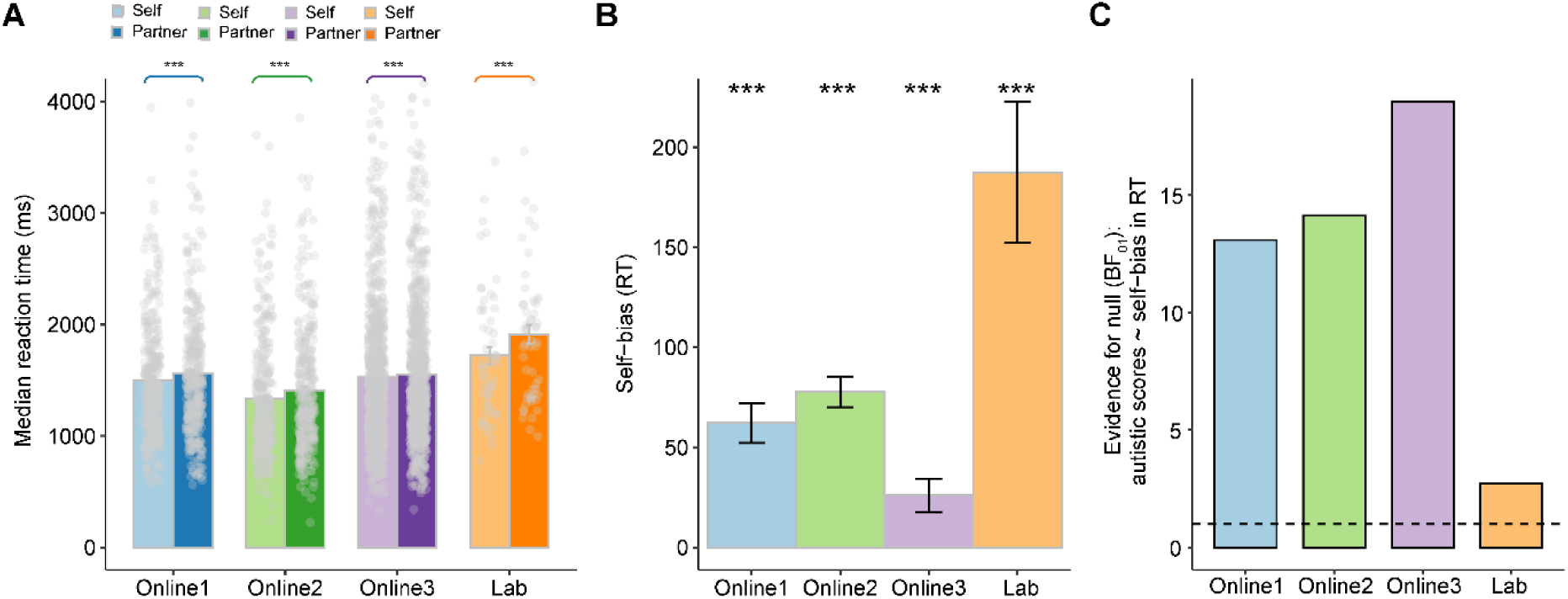
An alternative, independent measure of self-bias based on reaction times, further corroborated the absence of a correlation between self-bias and autistic trait score. **(A)** First, in all four studies, median reaction times provided a robust behavioural measure of the self-bias. Bright colors represent self-decisions and dark colors represent partner decisions in each experiment. In online experiment 1, participants responded quicker in self decisions compared to partner decisions (paired samples t-test: *t*(347) = −6.276; *p* < 0.001). Faster responses for self-related information can be considered as an alternative expression of the self-bias. Participants consistently responded quickly in self decisions compared to partner decisions across online experiment 2 (*t*(341) = −10.332; *p* < 0.001), online experiment 3 (*t*(874) = −3.131; *p* = 0.002), and lab experiment (*t*(55) = −5.299; *p* < 0.001), replicating the results observed in online experiment 1 (Error bars depict SEM. *** *p* < 0.001, ** *p* < 0.01, * *p* < 0.05.). **(B)** Self-bias in reaction time (RT) was quantified as the difference in median reaction time between partner and self decisions (self-bias in RT = RT_partner_ – RT_self_), where higher values reflect faster choice in self decisions. All bars are significantly positive as they quantify the effects shown in panel A in a single variable. **(C)** Correlations between self-bias in RT and autistic trait scores were tested across four experiments. Bayesian analyses provided strong evidence for the absence of a correlation across all studies (online experiment 1: BF_01_ = 13.09, online experiment 2: BF_01_ = 14.13, online experiment 3: BF_01_ = 18.98; lab experiment: BF_01_ = 2.708, with the combined dataset yielding BF_01_ = 27.27). The dashed line indicates the BF_01_ threshold of 1, above which values provide evidence in favour of the null hypothesis.

**Supplementary Figure S2.**
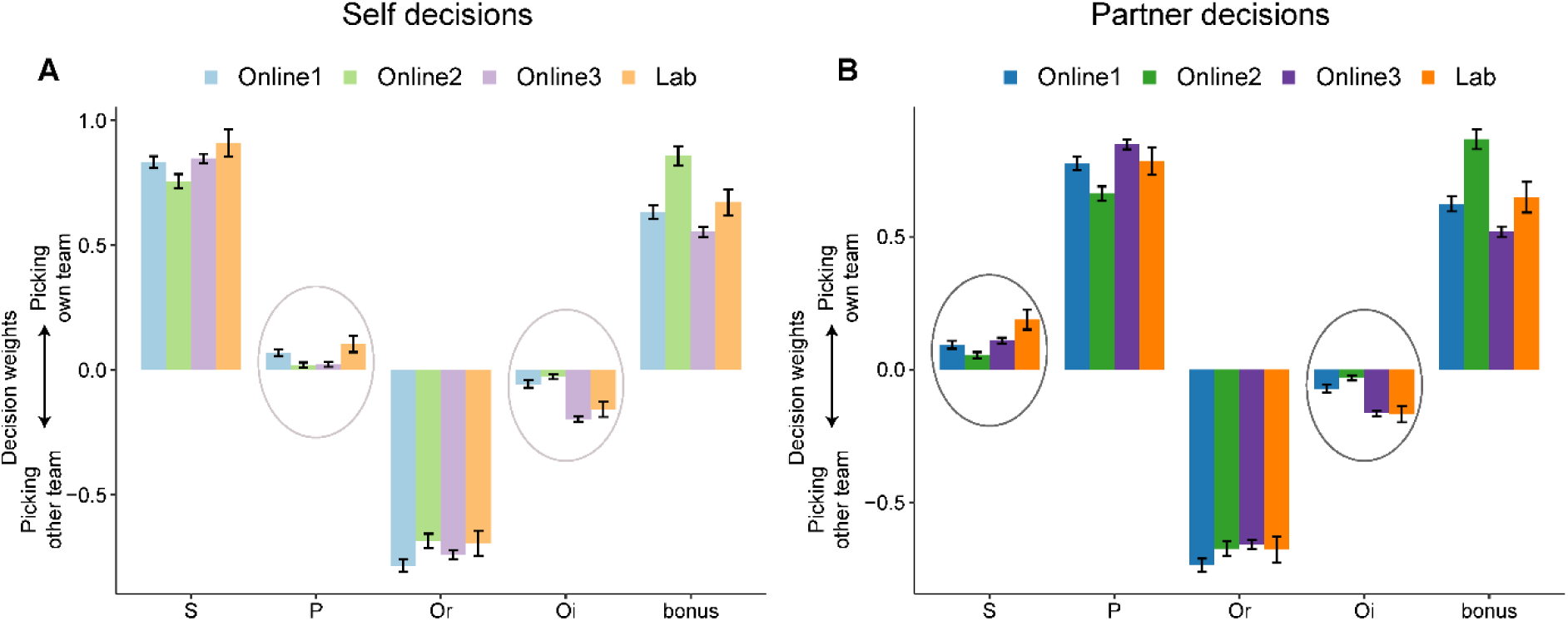
Full behavioural GLM results in self and partner decisions across four experiments. **(A)** In self decisions, participants’ decisions were mainly based on the performance scores from the self (S), the relevant opponent (Or), and the bonus. However, we also observed the small effects from the irrelevant players e.g., the partner (P) and irrelevant opponent (Oi), see the main text for the significance. Furthermore, these small effects were consistent with group membership: a better performance by P led to a larger likelihood of picking their team, whereas a higher score from Oi led to a higher probability of picking the opponent’s team. **(B)** In partner decisions, participants mainly made their decisions based on the performances of the partner and the relevant opponent. similar subtle effects from the irrelevant players were also observed across the four experiments. Error bars depict SEM.

**Supplementary Figure S3.**
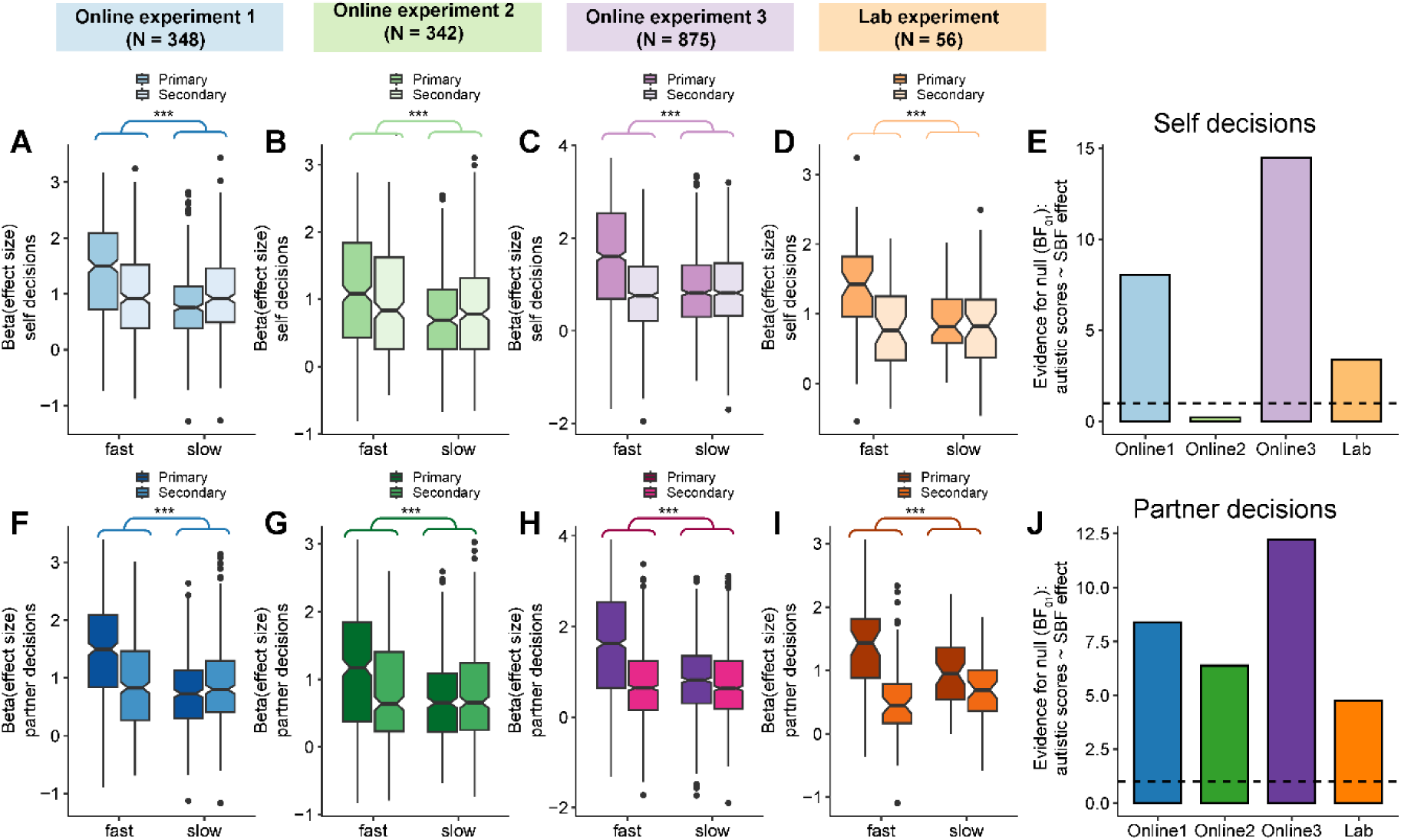
The absence of a correlation between social basis function use and autistic trait scores is further supported by an alternative signature of social basis function use. Previous work has shown that social basis function use is characterized by a step-wise decision making signature: people first identify the group structure (i.e., one team versus the other) via a primary basis function and then specify the relevant players in each team using a secondary basis function [1]. This model predicts that fast decisions are more strongly influenced by primary than secondary basis functions, the latter of which has not yet exerted its influence on choices. A lasso GLM (lambda = 0.012) was applied separately for decisions split into fast and slow trials by the median reaction time. We then predicted choice as a linear combination of primary and secondary basis function projection, and the bonus. An ANOVA was performed in the resulting beta coefficients of basis functions (primary, secondary) and reaction time (fast, slow) as within-subject factors, separately for self and partner decisions. **(A-D)** These results were consistent with our predictions and our previous work [1, 2] across all four studies: primary basis functions exerted stronger effects on choice than secondary basis functions in fast compared to slow decisions when decisions were made for self (online experiment 1: (*F*(1,347) = 122.1, *p* < 0.001; online experiment 2: (*F*(1,341) = 59.99, *p* < 0.001; online experiment 3: *F*(1,874) = 319.6, *p* < 0.001; lab experiment: *F*(1,55) = 17.40; *p* < 0.001). **(E)** Social basis function use was characterised by an earlier reliance on group structure than individual identity, with this effect being more pronounced in fast than slow decisions. We therefore used a quantification of social basis function use that is complementary to the analyses performed in the main manuscript. We calculated the difference in beta coefficients between primary and secondary basis functions in fast decisions, and then correlated this difference with autistic trait scores. In self decisions, Bayesian analyses provided strong evidence for the absence of correlation across studies, except online experiment 2 (online experiment 1: BF_01_ = 8.03, online experiment 2: BF_01_ = 0.204, online experiment 3: BF_01_ = 14.47; lab experiment: BF_01_ = 3.38, with a combined dataset supported the null: BF_01_ = 27.28). **(F-I)** In partner decisions, the same temporal prioritization on primary over secondary basis function, was replicated across four studies (online experiment 1: *F*(1,347) = 137.1, *p* < 0.001; online experiment 2: *F*(1,341) = 106.3, *p* < 0.001; online experiment 3: *F*(1, 874) = 272.3, *p* < 0.001; lab experiment: *F*(1,55) = 24.98, *p* < 0.001). **(J)** Similarly, no correlation was found between the alternative indice of social basis function use and autistic trait scores in partner decisions. Bayesian analyses provided strong evidence for the null across all four studies (online experiment 1: BF_01_ = 8.379, online experiment 2: BF_01_ = 6.382, online experiment 3: BF_01_ = 12.23; lab experiment: BF_01_ = 4.763; The combined dataset supported the null: BF_01_ = 31.04). The dashed line indicates the BF_01_ threshold of 1, above which values provide evidence in favour of the null hypothesis.

**Supplementary Figure S4.**
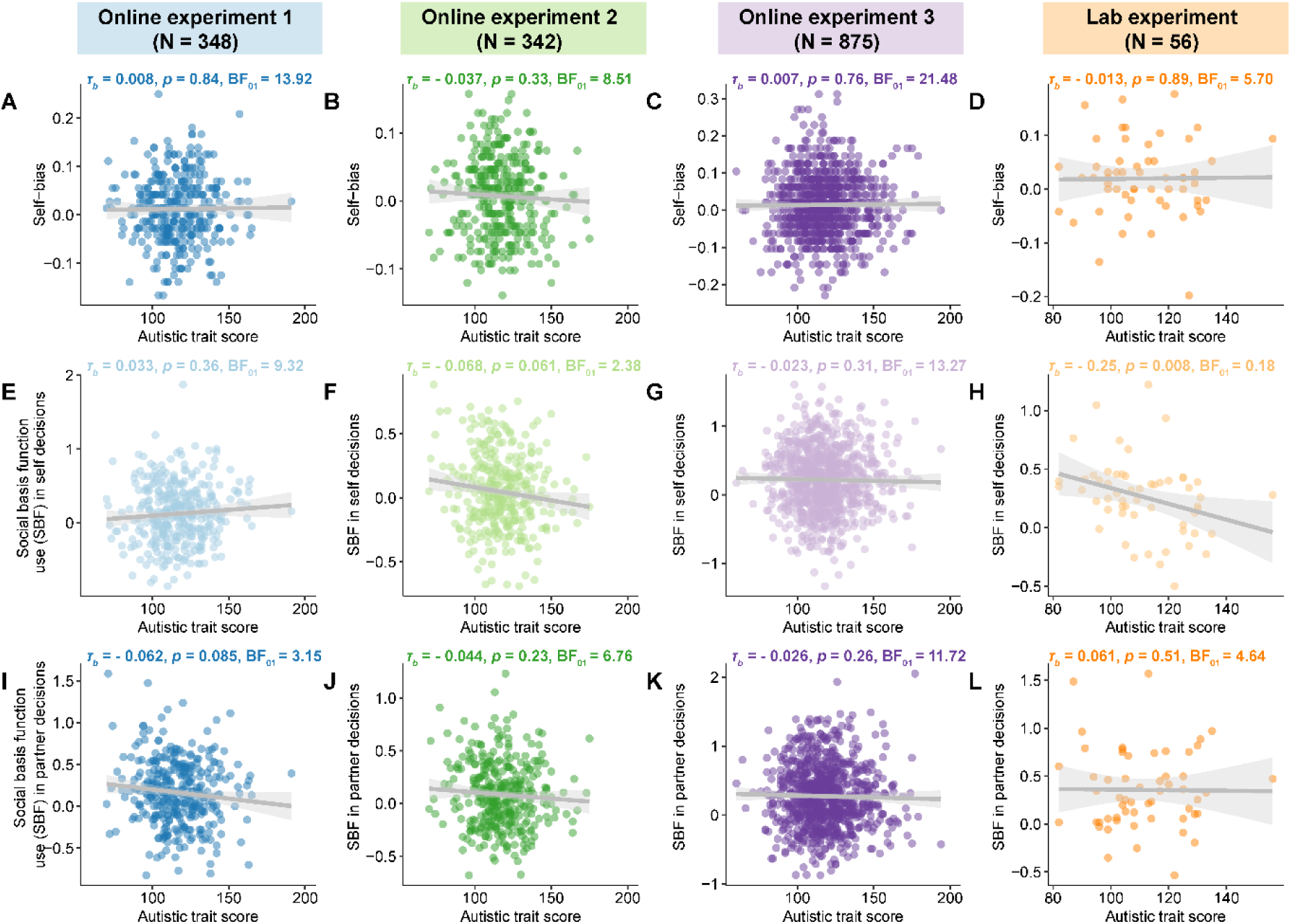
Kendall’s rank correlation between autistic trait score and self-other integration. We used Kendall’s rank correlation to reanalyse the relationship between autistic trait scores and self-other integration, which replicated all the non-significant results in Figure 4: self-bias (online experiment 1: BF_01_ = 13.92; online experiment 2: BF_01_ = 8.51; online experiment 3: BF_01_ = 21.48; lab experiment: BF_01_ = 5.70); social basis function use in self decisions (online experiment 1: BF_01_ = 9.32; online experiment 2: BF_01_ = 2.38; online experiment 3: BF_01_ = 13.27); social basis function use in partner decisions (online experiment 1: BF_01_ = 3.15; online experiment 2: BF_01_ = 6.76; online experiment 3: BF_01_ = 11.72; lab experiment: BF_01_ = 4.64). Note that the marginally significant correlation in online experiment 2 (self decisions, panel F) and online experiment 1 (partner decisions, panel I) became non-significant under Kendall’s rank analysis. Only in the lab experiment (panel H) did the correlation between social basis function use in self decisions and autistic trait scores remain significant (BF_01_ = 0.18). Overall, the results from Kendall’s rank correlation aligned with the findings presented in the main text.

**Supplementary Fig.S5.**
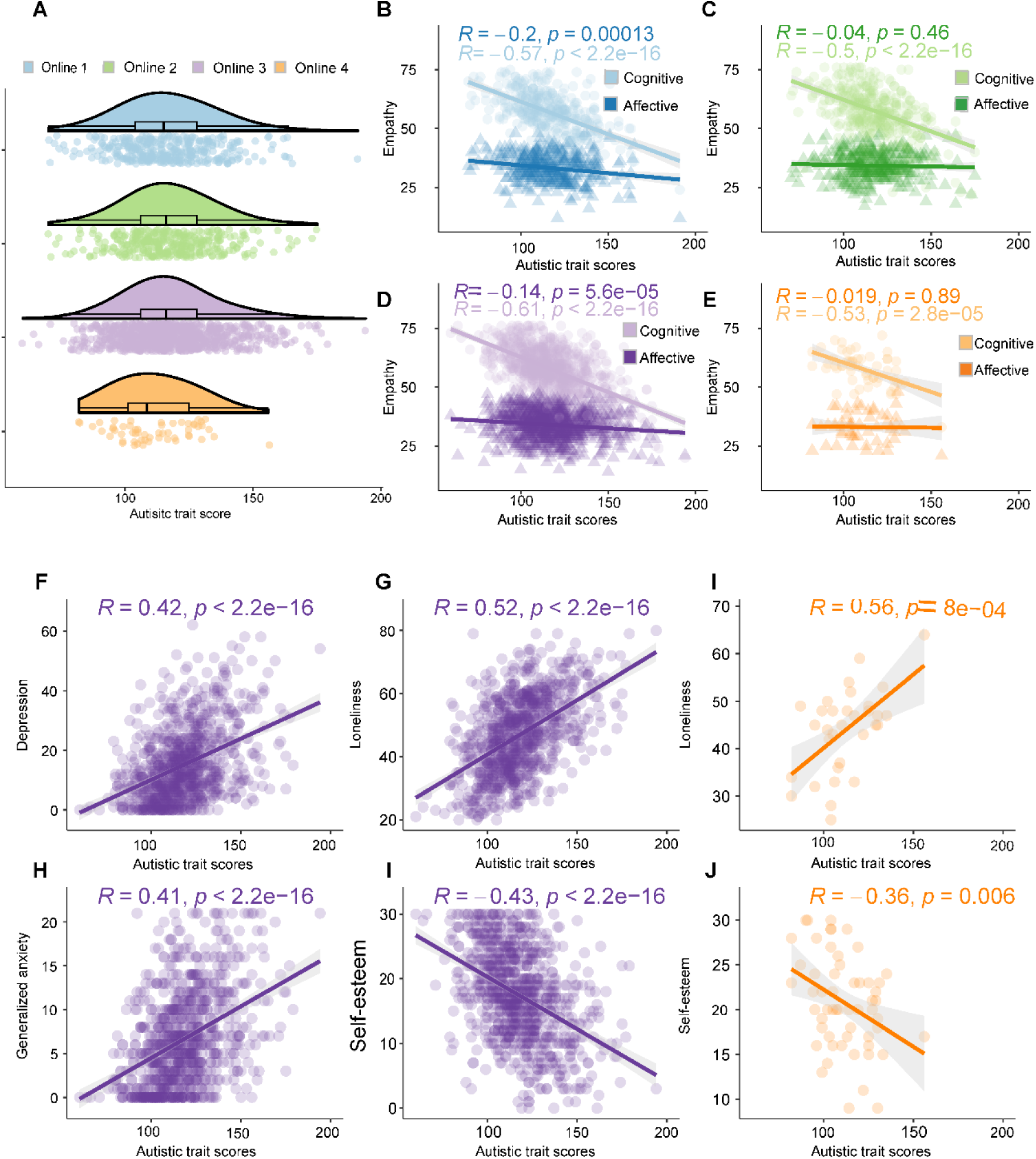
Expected patterns of correlation between autistic trait score, empathy, and mental health measures. **(A)** The distributions of autistic trait scores were consistent and similar across four experiments (*F*(3,1617) = 2.390, *p* = 0.067, η2 = 0.004); online experiment 1: ranged 70 to 191 (M = 116.6, SD = 18.31; online experiment 2: ranged 70 to 175 (M = 117.3, SD = 17.48); online experiment 3: ranged 60 to 194 (M = 117.7, SD = 18.52); lab experiment: ranged 82 to 156 (M = 111.3, SD = 15.1)). Note that the ANOVA is close to significant, and given reports that populations recruited online often score higher in mental health-related questionnaires [3, 4], we specifically considered whether the average AQ scores are higher in the online studies compared to the lab study. To do so, we used a Welch two-sample t-test, which accounts for the large difference in sample size between the lab study (n = 56) and the pooled online studies (n = 1565). Indeed, autistic trait scores were significantly increased in the online sample (*t*(60.92) = −2.964, *p* = 0.004). Note that this result also holds when calculating a standard Student’s t-test. Each color represents the data from one of the four experiments in all panels (online experiment 1: blue; online experiment 2: green; online experiment 3: purple; lab experiment: orange). Note that the set of questionnaires used varied across experiments. The following questionnaire correlations are illustrated in various colors corresponding to the data from the specific experiment. **(B, C, D, E)** Across all four experiments, autistic trait scores showed a stronger negative correlation with cognitive empathy (dark color) compared to affective empathy (light color) (online experiment 1: z = – 6.853, *p* < 0.001; online experiment 2: z = −7.814, *p* < 0.001; online experiment 3: z = −14.366, *p* < 0.001; lab experiment: z = −3.349, *p* < 0.001). This pattern aligns with previous findings that autism is linked to changes in the ability to take others’ perspectives, while still resonating with others emotionally in a similar manner as other people [5, 6]. **(F)** Autistic trait scores positively correlated with depression in online experiment 3. **(G, H)** Autistic trait scores positively correlated with loneliness in both online experiment 3 and the lab experiment (online experiment 3: r = 0.52, *p* < 0.001; lab experiment: r = 0.56, *p* < 0.001). **(I)** A positive correlation was observed between autistic trait scores and generalized anxiety in online experiment 3 (r = 0.41, *p* < 0.001). **(J, K)** Autistic trait scores negatively correlated with self-esteem in both online experiment 3 and the lab experiment (online experiment 3: r = - 0.43, *p* <0.001; lab experiment: r = - 0.36, *p* = 0.006). In summary, our measures of autistic trait score repeatedly and consistently show expected patterns of correlations with empathy as well as with measures of poor mental health.

**Supplementary Fig.S6.**
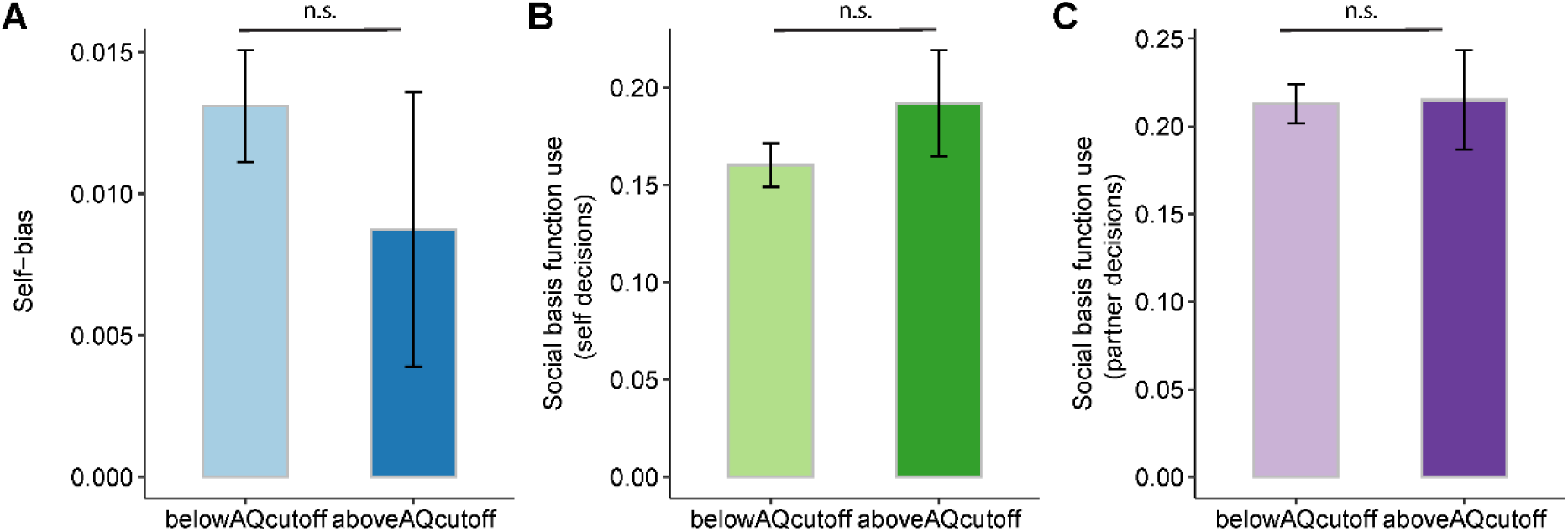
The markers of self-other integration unaffected in people scoring above the AQ clinical threshold. We combined datasets from all four experiments since we had demonstrated that these experiments showed similar task properties and expected questionnaire constructs, while also maximizing the sample size to achieve greater statistical power. We then catergorised participants based on the established clinical threshold [7–9], using an AQ score of ⩾ 32 with the binary scoring system. This identified 200 out of 1621 participants above the AQ cutoff across experiments. **(A)** Self-bias did not differ significantly between the group with autistic traits above the clinical threshold and those below the threshold (*t*(1619) = −0.778, *p* = 0.437, BF_01_ = 8.83). **(B)** Social basis function use in self decisions remained comparable across groups (*t*(1619) = 1.012, *p* = 0.312, BF_01_ = 7.19). **(C)** Likewise, the pattern of social basis function use in partner decisions aligned between the groups (*t*(1619) = 0.070, *p* = 0.945, BF_01_ = 11.84). These results also hold when analysing the online data sets exclusively, which we did because they had more comparable settings. The results also held when using two-sample Welch’s tests to account for the unequal sample sizes between participants below and above the cutoff. Overall, participants scoring above the threshold showed an unaltered pattern across all three measures of self-other integration (Error bars depict SEM; n.s.: *p* > 0.30.)

**Supplementary Fig.S7.**
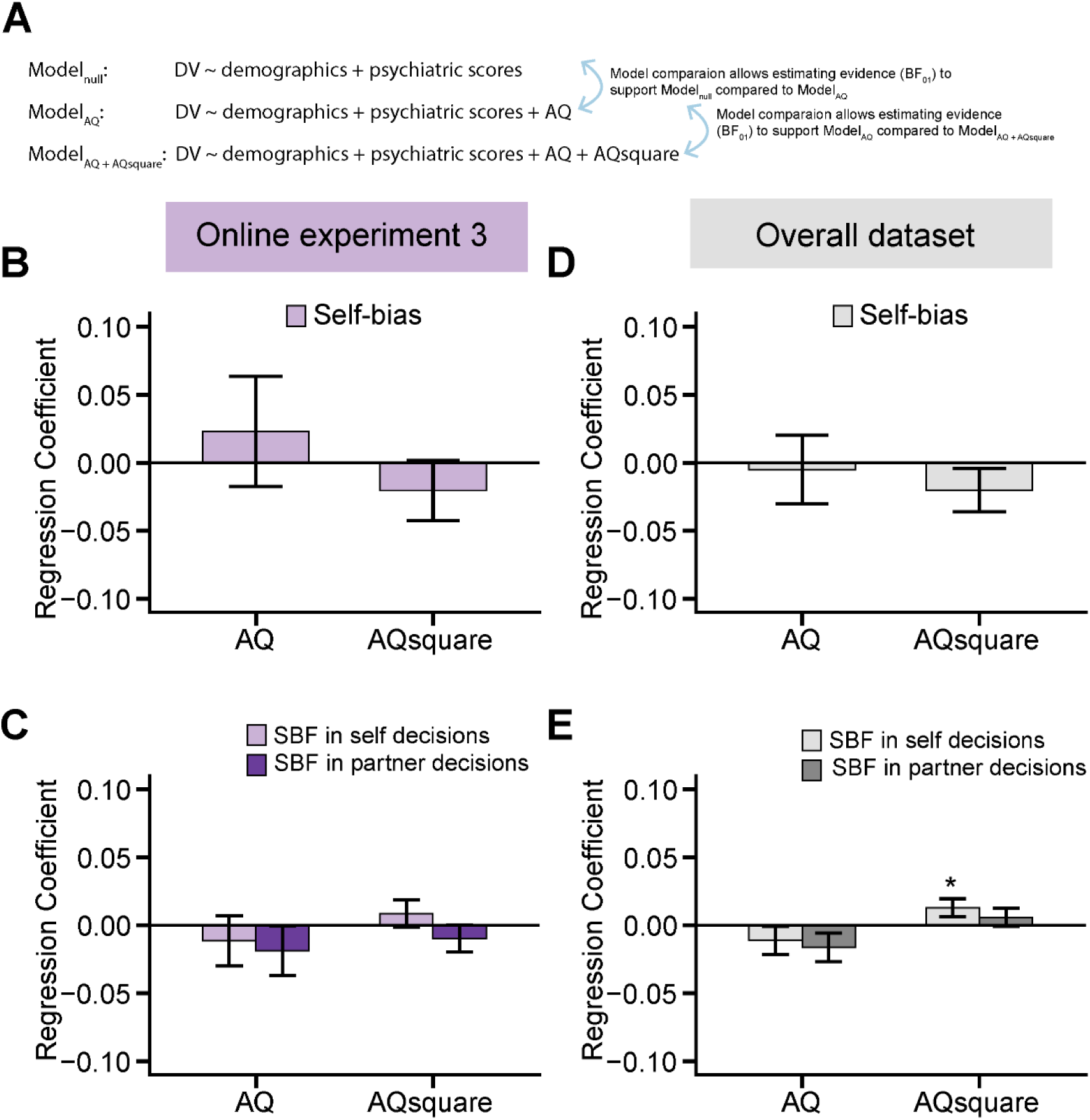
Relationship between autistic trait score and self-other integration is not masked by other markers of individual differences. **(A)** We have shown that self-other integration markers are unrelated to autistic trait score across our four experiments. Here, we sought to rule out that other markers of individual difference may have “masked” relationships between the AQ-50 and self-bias or social basis function use (SBF). We did this in two ways. We first re-ran our analysis, controlling for other individual difference factors. We constructed three general linear models that predicted our behavioural measures of self-other integration (self-bias, SBF in self and partner decisions) using a linear combination of demographic and questionnaire variables. The three models were nested inside each other. One variant of this model did not include the AQ score (model_null_), a second variant was identical to the first but now included the AQ score (model_AQ_), and a third model included in addition both the AQ score and the squared AQ score (model_AQ + AQ-square_). The last model allowed us to test whether our behavioural variables may exhibit a U-shaped relationship with autistic trait scores. U-shaped relationships are often found in mental health or developmental research [10–12]. We tested the beta weights of the AQ (in the model_AQ_) and AQ-square (in the model_AQ + AQ-square_) for significance, and in addition calculated the Bayesian model evidence favouring the null. **(B,C)** First, we examined online experiment 3 specifically, because it had the largest sample size (N=875) and comprised the highest number of control questionnaires (Fig.1 & Methods). This approach allowed us to test the correlations of interest while controlling for an extensive set of demographics and mental health variables. Our results revealed that neither autistic trait scores nor their squared terms predicted self-bias (|βs| ≤ 0.023, *ps* ≥ 0.356, panel B) or social basis function use in both self and partner decisions (|βs| ≤ 0.019, *ps* ≥ 0.305, panel C). Bayesian model comparison showed evidence in support of model_null_ as opposed to model_AQ_ (self-bias: BF_01_ = 3.42; social basis function use in self decisions: BF_01_ = 3.42; social basis function use in partner decisions: BF_01_ = 2.50). Similarly, Bayesian model comparison showed evidence in favour of model_AQ_ instead of model_AQ + AQ-square_ (self-bias: BF_01_ = 2.54; social basis function use in self decisions: BF_01_ = 2.77; social basis function use in partner decisions: BF10 = 2.47). **(D,E)** Second, we extended our analysis to include data aggregated across all four experiments (N = 1,621) to examine the relationship between autistic trait score and self-other integration, controlling for common potential confounds. We followed the same rationale as above comparing three general linear models. We found a consistent absence of relationships for both autistic trait scores and their squared terms in predicting self-bias (|βs| ≤ 0.020, *ps* ≥ 0.205; panel D) and social basis function use in partner decisions (|βs| ≤ 0.016, *ps* ≥ 0.119). We found no effect of autistic trait scores on social basis function use in self decisions (β = −0.011, *p* = 0.286). While their squared terms showed a subtle effect (β = 0.013, *p* = 0.048, panel E), this did not survive multiple comparison corrections. Bayesian model comparison showed evidence in support of model_null_ compared to model_AQ_ (self-bias: BF_01_ = 7.56; social basis function use in self decisions: BF_01_ = 4.47; social basis function use in partner decisions: BF_01_ = 2.39), and evidence in favour of model_AQ_ instead of model_AQ + AQ-square_ (self-bias: BF_01_ = 3.12; social basis function usein self decisions: BF_01_ = 1.03; social basis function usein partner decisions: BF_01_ = 4.80). Overall, our findings suggest absent significant relationships between autistic trait score and self bias, and to a lesser extent, no relationship between autistic trait score and social basis function use , after controlling for potential confounds.

## Notes

### Competing Interest Statement

The authors have declared no competing interest.

